# Integration of ethnobotany and population genetics uncovers the agrobiodiversity of date palms of Siwa Oasis (Egypt) and their importance to the evolutionary history of the species

**DOI:** 10.1101/820407

**Authors:** Muriel Gros-Balthazard, Vincent Battesti, Sarah Ivorra, Laure Paradis, Frédérique Aberlenc, Oumarou Zango, Salwa Zehdi, Souhila Moussouni, Summar Abbas Naqvi, Claire Newton, Jean-Frédéric Terral

## Abstract

Crop diversity is shaped by biological and social processes interacting at different spatiotemporal scales. Here we combined population genetics and ethnobotany to investigate date palm (*Phoenix dactylifera* L.) diversity in Siwa Oasis, Egypt. Based on interviews with farmers and observation of practices in the field, we collected 149 date palms from Siwa Oasis and 27 uncultivated date palms from abandoned oases in the surrounding desert. Using genotyping data from 18 nuclear and plastid microsatellite loci, we confirmed that some named types each constitute a clonal line, i.e. a true-to-type cultivar. We also found that others are collections of clonal lines, i.e. ethnovarieties, or even unrelated samples, i.e. local categories. This alters current assessments of agrobiodiversity, which are visibly underestimated, and uncovers the impact of low-intensity, but highly effective, farming practices on biodiversity. These hardly observable practices, hypothesized by ethnographic survey and confirmed by genetic analysis, are enabled by the way Isiwans conceive and classify living beings in their oasis, which do not quite match the way biologists do: a classic disparity of *etic* vs. *emic* categorizations. In addition, we established that Siwa date palms represent a unique and highly diverse genetic cluster, rather than a subset of North African and Middle Eastern palm diversity. As previously shown, North African date palms display evidence of introgression by the wild relative *Phoenix theophrasti*, and we found that the uncultivated date palms from the abandoned oases share even more alleles with this species than cultivated palms in this region. The study of Siwa date palms could hence be a key to the understanding of date palm diversification in North Africa. Integration of ethnography and population genetics promoted the understanding of the interplay between diversity management in the oasis (short-time scale), and the origins and dynamic of diversity through domestication and diversification (long-time scale).

## 1. Introduction

The date palm, *Phoenix dactylifera* L., is a major perennial crop of the hot and arid regions in the Middle East and North Africa (Barrow, 1998). Its sugar-rich fruit, the date, has been consumed for millennia (Tengberg, 2012), and has long been rooted in Berber/Amazigh and Arabic cultures. *Phoenix dactylifera* belongs to the Arecaceae family and, along with 12 or 13 other interfertile species, composes the genus *Phoenix* (Barrow, 1998). Date palms exist mostly as either cultivated or feral (i.e. uncultivated but derived from cultivated palms) domesticated forms (for review, Gros-Balthazard et al., 2018). Only a few relictual populations of its wild progenitor are known today in Oman (Gros-Balthazard et al., 2017), even though Tuaregs of the Tassili n’Ajjer (Algeria) consider it to be wild in their gardens (Battesti, 2004). *Phoenix dactylifera* is dioecious, meaning that an individual date palm is either male or female, and, naturally, only females bear fruit. Today, thousands of female cultivars are reported, varying in fruit shape, color, texture or taste, but also in their vegetative aerial architecture (Chao and Krueger, 2007). Without addressing the existence of male “cultivars” (there is little or no vegetative reproduction of identified and named males except in research stations), male varieties are also locally identified, but are less studied. Further, local categorization by farmers requires clarification, starting with the adequacy of the notion of “cultivar” or what it is supposed to be.

In palm gardens, palm trees bear a name. The unquestioned and often implicit assumption is that each female date palm named type (for instance the famed Medjool or Khalas) is a cultivar, that is a clone, multiplied only through vegetative propagation. Phoeniculture, or date palm cultivation, involves a mix of clonal and sexual propagation. But in order to obtain specifically female trees and to ascertain that fruits will be of the desired, predictable quality, farmers mostly make use of the asexual reproduction abilities of this plant through offshoot multiplication. Indeed, sexual reproduction of date palms only leads in rare cases to progenies having equivalent or superior fruit qualities (4‰ according to Peyron, 2000), although these assessments remain subjective. For the oasis system to be efficient in the Sahara, where the scarcity of water, irrigable lands, and manure is a real concern, all oasis communities chose the option of planting and maintaining 95-99% of female palms (Battesti, 2005) instead of a natural 50:50 sex ratio (Chao and Krueger, 2007). Opting for such an artificial sex ratio requires hand-pollination, a practice already used in southern Mesopotamia during the 3^rd^ millennium BC (Landsberger, 1967), as a substitute to natural wind pollination (Henderson, 1986). Hence, farmers almost entirely restrict sexual reproduction by seeds, with a few exceptions (e.g. in India, Newton et al., 2013). Accidental seedlings are however sometimes spared. A resulting male may be later used for pollination. If female, its fruits are sometimes harvested and, although rare, it can lead to a new selected and named line of clones i.e. a cultivar. Another possibility, highlighted in our previous work in the oasis of Siwa, is to integrate this new genotype into an existing named type, because, from a local perspective, it is the very same variety, the same “form” or phenotype (Battesti, 2013; Battesti & Gros-Balthazard et al., 2018). This farming practice challenges the presumption of a named type being a true-to-type cultivar, i.e. aggregating solely vegetatively propagated individuals. This may, in turn, lead to a misinterpretation and an inaccurate estimation of local agrobiodiversity.

Siwa is a desert oasis located in the Libyan Desert 300 km south of the Mediterranean coast and the closest city Marsa Matruh, and about 30 km east of the Libyan border (Figure 1, Figure S1A). It is located at the breaking of the limestone Miocene plateau at the edge of the Qattara Depression and at the northern limit of the Great Sand Sea. Much of the depression sits below sea level. A large part of this territory is occupied by salt lakes, “*sebkha*”, largely resulting from agricultural activity (significant drainage is provided to leach the soil and protect it from salinization). The cultivated area extends in the form of palm groves over the rest of the territory, but is highly concentrated around settlement areas (Figure 1). With roughly 200,000 to 250,000 palms (Battesti, 2013), dates are the main commercial crop in Siwa Oasis, closely followed by olive (*Olea europaea* L.). As in most oasis systems, date palm is used to feed the oasis inhabitants, but also for drinks, fodder, building materials (beams, hedges), crafts (baskets, various utensils), and daily uses (ropes, ties, brooms, furniture) (Battesti, 2005). The date palm is of primary importance, more generally, as the keystone of the oasis ecological system (microclimate effect, Riou, 1990). It is also the cornerstone of the local economy of exportation to large urban centers. The oases have a very relative autarchy and self-sufficiency, their inhabitants export what they have in abundance: dates (Battesti, 2013; Battesti & Gros-Balthazard, et al., 2018). Extrapolating from early 20^th^ century data in Siwa, an estimate of 82% of the dates in value terms were exported (and 95% of the elite cultivar □a□idi) (Hohler and Maspero, 1990). Several population genetics analyses focused on date palm agrobiodiversity in Siwa (Abd El-Azeem et al., 2011; Abou Gabal et al., 2006; El-Sharabasy and Rizk, 2019; el-Wakil and Harhash, 1998; Hemeid et al., 2007; Selim et al., 1970). Nevertheless, none of these studies integrate agricultural knowledge and practices, and the existing diversity in Siwa Oasis and its categorization system remains thus poorly understood.

**Figure 1.**
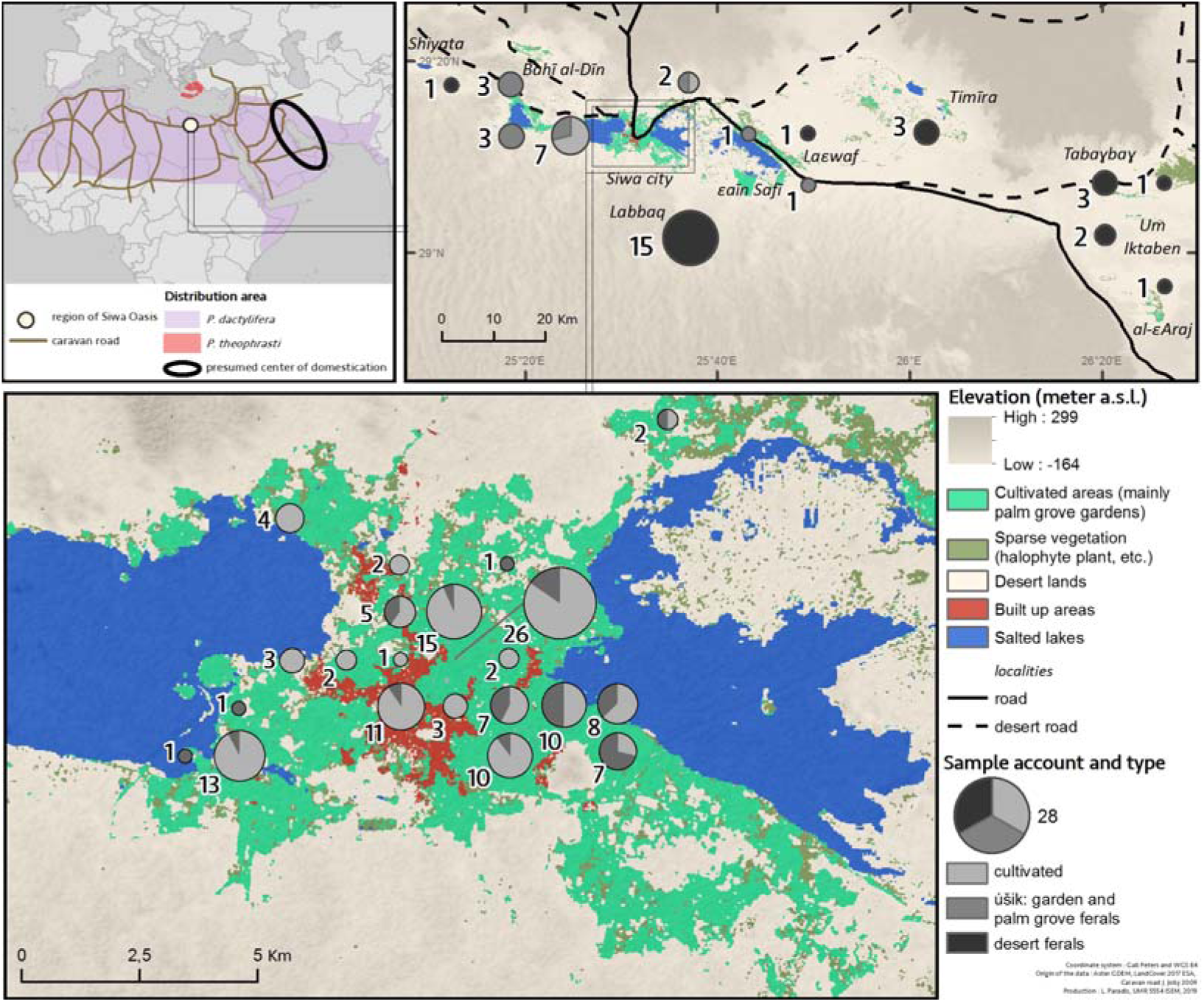
Localization of Siwa Oasis and sampling strategy in Siwa region.

In Siwa Oasis, we previously described a complex system where named types are not necessarily cloned genotypes, but so-called ethnovarieties or local categories (Battesti, 2013; Battesti & Gros-Balthazard, et al., 2018). To synthesize our “objectification” of the local organization of the date palm agrobiodiversity system, we proposed these definitions:

- *Cultivar or true-to-type cultivar*: a set of clonal individuals, i.e., the association of a name and a single genotype reproduced vegetatively (asexually, by offshoot) by humans. Genotypes of a same cultivar are genetically identical, except in case of somatic mutations (McKey et al., 2010).

- *Ethnovariety*: a set of similar (according to local standards) lines of clonal individuals reproduced vegetatively (asexually, by offshoot) by humans deliberately under a single local name.

For instance, our ethnobotanical survey pinpointed that the two named types úšik niqbel and □rom a□za□1 were presumably true-to-type cultivars i.e. to only come from offshoots and never possibly from a seedling (Battesti, 2013). Nevertheless, our genetic analysis indicated that ús□ik niqbel was more likely a collection of multiple clones, a.k.a. an ethnovariety, rather than a true cultivar, while we did not have enough data to rule on □rom a□za̅1 (Battesti, Gros-Balthazard, et al., 2018).

We also hypothesized that the named types might meet another order of categorization: inclusive local categories, the most obvious of which are male (*ó□em*) and female seedlings (úšik). We can therefore add this definition:

- *Local category*: a set of individuals sharing some characteristics (according to local standards i.e. fruit color, harvesting season, usage, rusticity), typically found as seedlings, sometimes reproduced vegetatively (asexually, by offshoot) by humans but identified under a single local name.

Indeed, a given name can also refer to an inclusive local folk category of date palms. So, the two largest “local categories” are the *ótem*, the male date palms, and *úšik* (not to be confused with the named types with the form *úšik xxx*, as *úšik n gubel*), which includes all date palms resulting from sexual reproduction (a seedling and not an offshoot). Both are said to have “no (further specific) names”, even if the farmers acknowledge a high degree of heterogeneity, and can “qualify” them individually: as *úšik ma asil*, whose dates let honey drip out, or *úšik □alaf*, whose dates are used as fodder, etc. As such, they are not conceived as varieties, but qualities. Our former results based on genetic data undeniably validated this hypothesis for *úšik*, but it was more puzzling for given names like ús□ik ezzuwa□ for instance (Battesti & Gros-Balthazard et al., 2018). The ethnographic survey pointed out the controversial status of this named type (is it a single clone or not) among farmers in Siwa Oasis (Battesti, 2013). With genetic data, we finally proved it is not a clone (a true-to-type cultivar). Is it even an ethnovariety or just a local category, i.e. an ús□*ik* with a morphological specificity: round and reddish/dark dates (*zuwa* being related to the reddish/dark color in Amazigh language)?

The origins of this date palm heritage and Siwa Oasis altogether are lost in the mists of time. Siwa Oasis was well known during Antiquity, in its Egyptian dynastic period, under the name of “Se*x*et-□m”, the land of date palms (Duemichen, 1877), that included the current oasis but also the oases now abandoned in its periphery (Kuhlmann, 2013). In the classical period, the oasis was famous throughout the Mediterranean Basin for its oracle built in the 6^th^ century BCE (Kuhlmann, 1988, 2011; Leclant, 1950), under the name of Amon Oasis (later Hellenized in Ammon). Alexander the Great was one of the most famous to consult this desert oracle, for him to confirm his divine ascendancy before his campaign of conquest in Persia. The dates of Siwa were mentioned as early as the 5^th^ century BCE by Hellanicus of Mytilene (c. 480-c. 395 BCE) in his *Journey to the Oracle of Ammon* (cited by Leclant, 1950, p. 248). The date palms of Siwa were then mentioned or even celebrated by Theophrastus (c. 371-c. 288 BCE) and later Pliny the Elder (23-79 CE) and Arrian (c. 95-v. 175), before a long silence and a return under the famous Arabic authors al-Bakrī (1040-1094), al-Idrīsī (1100-1165), and al-Maqrīzī (1364-1442). A few genetic studies involved date palms from Siwa (Abd El-Azeem et al., 2011; Abou Gabal et al., 2006; Gros-Balthazard et al., 2017; Hemeid et al., 2007). Nevertheless, not all named types of the oasis were studied, levels of categorization of date palm names were confused (sometimes mistaking *úšik*, seedlings, for a cultivar), farming practices were neglected, and only date palms cultivated in the current oasis were considered, ignoring the abandoned oases scattered in the desert. A proper assessment of date palm agrobiodiversity in Siwa region, and in a broader sense an understanding of its origin, hence are still lacking.

Beyond the peculiar history of Siwa Oasis, the scenarios of the beginning of date palm cultivation in Egypt in particular, and in North Africa in general, remain incomplete. Older evidence of exploitation are found around the Persian Gulf while it seems that in North Africa, cultivation is more recent (Flowers et al., 2019; Tengberg, 2012). In Egypt, the date palm seems exploited or cultivated sporadically since the Old Kingdom (about 2700-2200 BCE), but phoeniculture is only established since the New Kingdom, about 1600-1100 BCE (Tengberg and Newton, 2016). Genetic analyses of the current date palm germplasm identified two differentiated genetic clusters in North Africa and the Middle East, with evidence of gene flows, especially in Egypt (Flowers et al., 2019; Gros-Balthazard et al., 2017; Hazzouri & Flowers, et al., 2015; Zehdi-Azouzi et al., 2015). A recent study showed that the cultivated North African pool has mixed ancestry from Middle Eastern date palms and the Aegean endemic wild relative *Phoenix theophrasti* Greuter, a.k.a. the Cretan date palm (Flowers et al., 2019). Nevertheless, the geographic, chronological and historical contexts of this introgression remain enigmatic. The oasis of Siwa is located at the crossroads between Greek, Libyan and Egyptian influences. It is on one of the rare passage points, a rare node in the network (Battesti & Gros-Balthazard, et al., 2018), between the east and west of the distribution area of the cultivated date palm (Figure 1). A deep understanding of date palm diversity in the region could therefore enlighten the diversification history of *Phoenix dactylifera* in North Africa.

In this paper, we are taking our previous work on Siwa date palms (Battesti et al., 2018) to the next level, using a joint ethnographic study and genetic analysis. First, we greatly expanded the number of named types sampled in Siwa to further test whether named types are true-to-type cultivars or ethnovarieties or local categories. Secondly, we also increased our sampling of uncultivated individuals from the abandoned oases in the desert nearby Siwa (also known as “feral” in Battesti, Gros-Balthazard, et al., 2018). This enabled a full assessment of local biodiversity and potential connections between the currently cultivated pool of the oasis and the uncultivated, abandoned date palms from the surrounding desert. Lastly, we used genotyping data from more than 200 cultivated date palms sampled across the entire historical range of the species in order to locate Siwa diversity located within a wider germplasm diversity. By including *Phoenix theophrasti*, we can assess a possible gene flow in this particular population of date palms. From a broader perspective, our work aims at documenting the origins of phoeniculture in Egypt and in North Africa in general.

## 2. Material & Methods

### 2.1. Plant samples and genotyping

#### 2.1.1. Ethnobotanical study and sample collection

We sampled 176 cultivated and uncultivated date palms in Siwa Oasis (Egypt) and surrounding desert (Figure 1, Table 1, Table S1), of which 52 were included in our previous study on the agrobiodiversity of date palms in Siwa (Battesti & Gros-Balthazard, et al., 2018). Those samples were collected *in situ* with the essential cooperation of the local famers while conducting an ethnobotanical study (Figure S1B). VB conducted social anthropological fieldwork between 2002 and 2017, including about six months dedicated to date palm categorization and naming (Battesti, 2013). While he occasionally conducted structured interviews or held focus group discussions and free listings of date palm given names, most of his data derived from participant observation (Battesti et al., 2018). This collection is composed of 109 accessions of date palms growing in about 46 private gardens that were deliberately chosen scattered throughout the current oasis. For each named type, we collected more than one date palm, when possible (some are rare), in order to test, with genetic data, whether they represent cloned accessions or not. We also sampled accidental seedlings growing in gardens (referred to as ús□ik #1), and on the border of gardens or palm groves (referred to as ús□ik #2). Further, 27 uncultivated date palms growing in abandoned oases in the desert were sampled (Figure 1). Those oases probably already existed during the Roman/Ptolemaic period, or at least some of them, and have been presumably abandoned since the 9^th^ or 10^th^ centuries CE (Battesti, 2013).

**Table 1.**
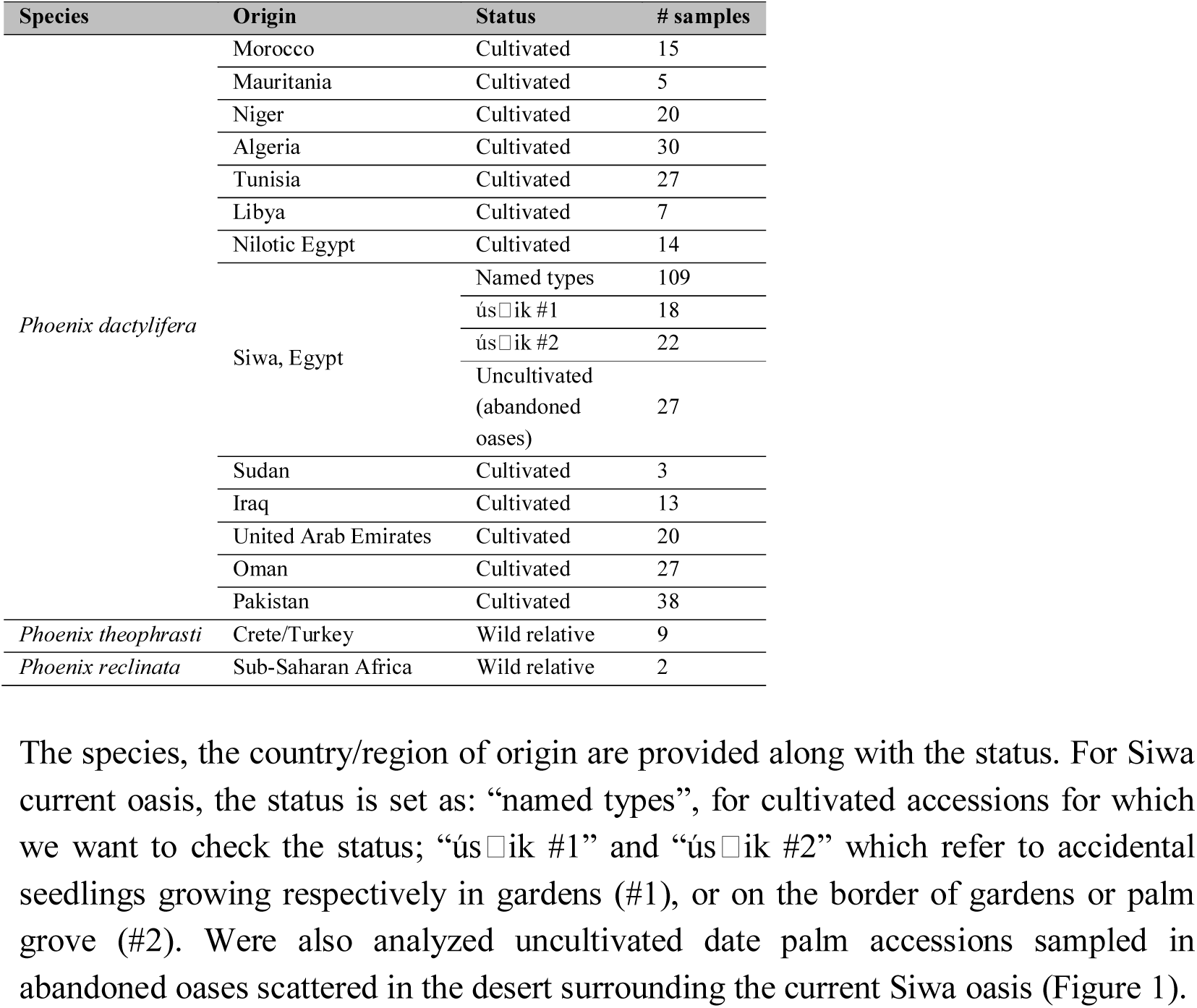
Summary of the 406 accessions of *Phoenix* spp. included in this study.

We additionally sampled nine *Phoenix theophrasti* Greuter in their native habitat in Crete and Turkey. Two accessions of *Phoenix reclinata* Jacq., collected in the botanical garden of the Villa Thuret in Antibes, France and originally from sub-Saharan Africa, were included as outgroup population. For each accession, a few leaflets were collected and dried either in the sun or by silica gel.

#### 2.1.2. DNA extraction and microsatellite genotyping

The collection was genotyped using 17 nuclear microsatellites and one chloroplastic minisatellite (Table 2), following the protocol of Zehdi-Azouzi et al. (2015). We crushed 40 mg of silica-dried leaves in a fine powder using bead-mill homogenizer TissueLyser (Qiagen, Courtabœuf, France). Total genomic DNA was extracted from leaf powder using DNeasy plant MINI Kit (Qiagen, Courtabœuf, France). The plastid dodecanucleotide minisatellite identified in the intergenic spacer psbZ-trnfM (Henderson et al., 2006) was genotyped. It has been previously used to define the date palm chlorotype (so-called occidental or oriental), depending on its number of repeats (three or four, respectively), and to barcode *Phoenix* species (Ballardini et al., 2013; Gros-Balthazard et al., 2017; Pintaud et al., 2010; Zehdi-Azouzi et al., 2015).

**Table 2.**
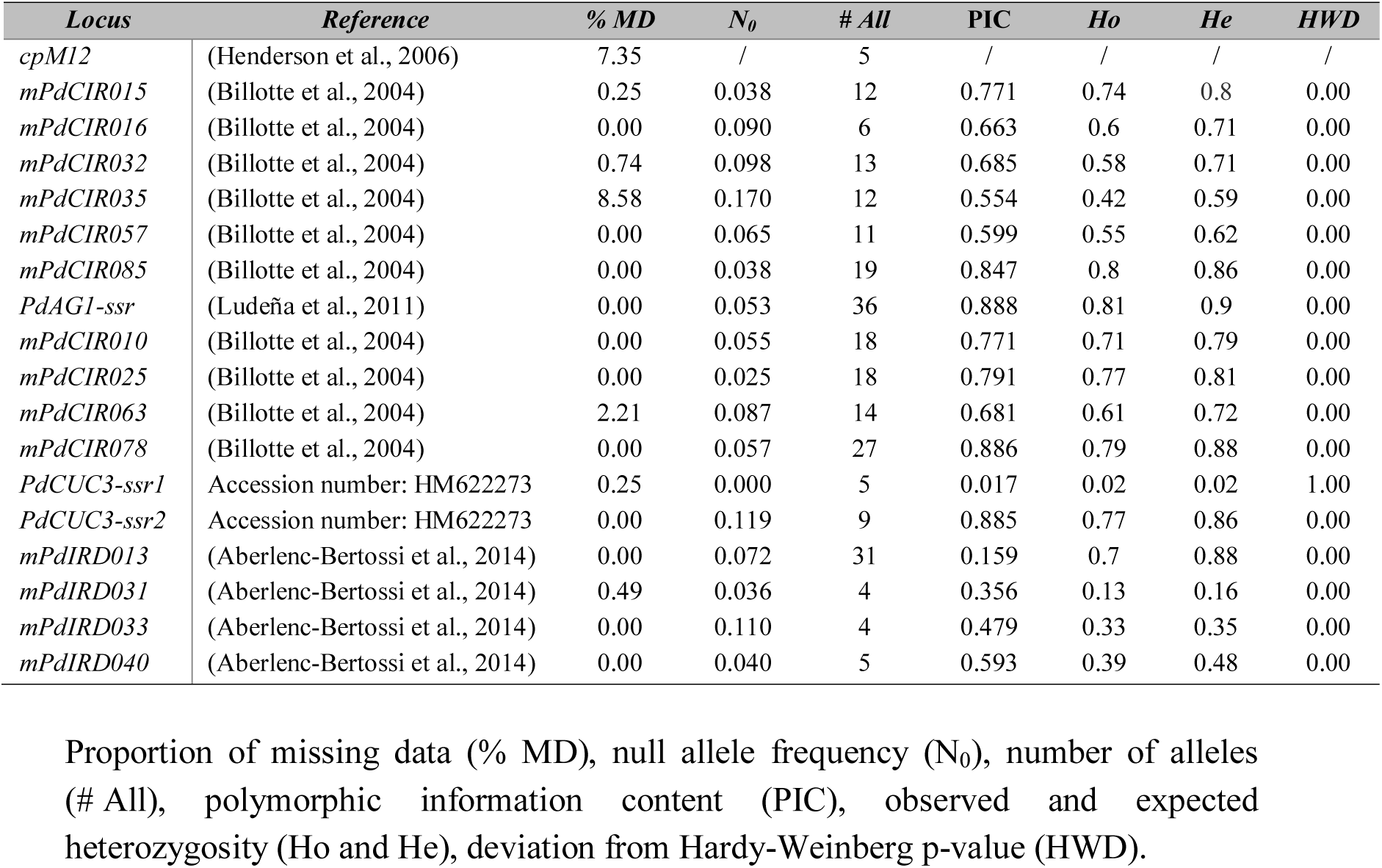
List of microsatellite loci and their summary statistics calculated on the 406 *Phoenix* spp. accessions included in the present study.

In addition to this newly generated genotyping dataset, we utilized *P. dactylifera* genotyping data from previous studies (Moussouni *et al*., 2017; Zango *et al*., 2017; Zehdi-Azouzi *et al*., 2015) (Table S1). Both this published data and our new data relied on the same set of microsatellite markers, and were generated by the same company (ADNid, Montpellier, France) with the same protocol, enabling a meta-analysis. These additional accessions are a good representation of the cultivated germplasm as their origin spans the historic date palm distribution, stretching from North Africa to the Middle East and Pakistan (Barrow, 1998).

### 2.2. Genotyping data analysis

Statistical analysis was conducted with the *R* Statistical Programing Language (R Core Team, 2015), unless otherwise stated. To identify duplicated genotypes among the whole dataset, we performed an identity analysis using Cervus v3.0.7 (Kalinowski et al., 2007). For each pair of accessions, Cervus calculates the number of matching genotypes across the 17 nuclear loci. We used the same software for calculation of the polymorphic information content (PIC) for each locus. We estimated null allele frequencies using *null.all* function in the *R* package *PopGenReport* (Adamack and Gruber, 2014; Gruber and Adamack, 2015). For each locus, the number of alleles N_A_, the observed (H_O_) and expected heterozygosity (H_S_) were estimated using the *R* package *pegas* (Paradis, 2010). The deviation from Hardy-Weinberg equilibrium was estimated using the function *hw.test* from the same package.

### 2.3. Date palm agrobiodiversity in Siwa

For each named type of Siwa, we checked whether they are actual true-to-type cultivars or rather represent a group of more or less distant genotypes (ethnovariety, local category). For this purpose, we calculated a measure of identity by averaging over each named type the proportion of matching genotypes across the 18 chloroplastic and nuclear loci. For true-to-type cultivars, this number is expected to be 100%, except in case of somatic mutations or genotyping error. Second, we calculated the Euclidean distance between each of the Siwa samples (function *dist*, *R* package *stats*) and built a heatmap of those distances using *heatmap* function (*R* package *stats*).

To investigate the extent of the diversity in the whole region, and not only in the oasis, we included uncultivated date palms from the abandoned oases in the following analyses. We generated a Neighbor-joining tree based on Nei’s genetic distance (Nei, 1972) using *aboot* function in *poppr* package (Kamvar et al., 2014) with 100 bootstrap replicates.

For each line of clones identified with these analyses, we kept a single accession for downstream analyses and performed a Principal Component Analysis using function *dudi.pca* in the *ade4* package (Dray and Dufour, 2007). Missing data were replaced by the mean allele frequencies using the function *scaleGen* from the *R* package *adegenet* (Jombart and Ahmed, 2011).

### 2.4. Extent and partitioning of worldwide date palm diversity

To draw up a picture of the overall *Phoenix* population structure, we applied two different approaches to our dataset comprising 128 unique genotypes from Siwa, 219 date palms originating from all over its historical distribution area, and nine wild *Phoenix theophrasti*. First, we performed a Principal Component Analysis, as described above for Siwa accessions only. We further used the Bayesian clustering method based on a Markov-chain Monte Carlo (MCMC) algorithm implemented in Structure v2.3.3 (Pritchard et al., 2000). Individuals are partitioned into a predefined number of clusters (*K*) so as to minimize linkage disequilibrium and deviation from Hardy-Weinberg within cluster. Allelic frequencies at each locus are calculated at the same time for each cluster. Genotyped individuals were allocated to one to eight clusters *K*. All runs were performed using a model allowing admixture and correlated allele frequencies among populations (Falush et al., 2003). A 100,000 iterations burn-in period was followed by 1,000,000 MCMC steps. Ten independent runs were performed for each specified *K*, the convergence of likelihood values was checked for each *K*. The optimal value of *K* was estimated using both the approach of Pritchard *et al*., 2000 based on the maximization of the log likelihood, and the approach of Evanno et al., 2005 based on the rate of change in the log likelihood between successive *K* values (delta *K*).

For each population pairs, we estimated a measure of differentiation (F_ST_) using the Genepop software v. 4.7 (Rousset, 2008). The proportion of shared alleles between populations and subpopulations was calculated with the *pairwise.propShared* function of the *R* package *PopGenReport* (Adamack and Gruber, 2014; Gruber and Adamack, 2015). Further, we calculated various diversity estimates for each population. The allelic richness and private allelic richness were calculated with a custom *R* script. Because the numbers of distinct alleles and private alleles depend heavily on sample size in each population, we used the rarefaction method (Petit et al., 1998), allowing a direct comparison among populations of different sample size. For each set of comparison, we thus used the smallest haploid sample size (n = 18 in *Phoenix theophrasti*) as the number of alleles to sample, and ran 1,000 replicates of allelic and private allelic richness calculation. To test for significant differences among populations, these diversity estimates were assigned to Tukey groups (function *HSD*.*test*, *Agricolae* package, de Mendiburu, 2015) at a significant threshold of 0.05. Expected and observed heterozygosity (H_S_ and H_O_) were calculated using the *basic.stats* function of the *R* package *hierfstat*. The inbreeding coefficient F_IS_ was calculated with the same package and confidence intervals were estimated by performing 1,000 bootstrap replicates over loci with the function *boot.ppfis*.

## 3. Results

A total of 406 *Phoenix* spp. samples (Table 1, Table S1) genotyped across 17 nuclear microsatellites and one chloroplastic minisatellite (Table 2) were analyzed in the present study. We report new genotyping data for 176 cultivated and uncultivated date palms from the Egyptian oasis of Siwa and the surrounding abandoned oases (Figure 1, Figure S1C), nine *Phoenix theophrasti*, and two *Phoenix reclinata* (Table S2). Additional genotyping data for 98 Middle Eastern/Asian and 121 North African date palms were retrieved from previous studies (Moussouni *et al*., 2017; Zango *et al*., 2017; Zehdi-Azouzi *et al*., 2015).

Missing data across the full dataset were very limited with an average of 1.1%, and the mean null allele frequency was 6.8%, on average across all 18 chloroplastic and nuclear loci (Table 2). All loci were polymorphic, with four to 31 alleles, and an average polymorphic information content of 0.625. All loci deviated significantly (*P* < 0.05) from Hardy-Weinberg equilibrium except Cuc3-ssr1. Except for some Siwa samples (see below), all accessions were unique, as no pair of accessions displayed 18 matches across the 18 chloroplastic and nuclear loci (Table S3).

**Table 3.** Named type of Siwa, sampling effort and genetic identity.

| Named type | # sample | Nuc Identity | Cp Identity |
| --- | --- | --- | --- |
| sa□idi [tasutet] | 13 | 97.59% | 100% |
| alkak | 11 | 100% | 100% |
| a□za□l | 4 | 97.06% | 100% |
| halu en □anem | 3 | 100% | 100% |
| tattagt | 8 | 100%* | 100% |
| ús□ik niqbel / ús□ik en gubel | 12 | 69.96% | 92.86% |
| □rom a□za□l | 6 | 28.02% | 88.03% |
| alkak wen z□emb | 6 | 28.24% | 88.44% |
| □rom sa□id | 2 | 29.41% | 0% |
| lekrawmet | 4 | 40.20% | 100% |
| ús□ik ama□zuz/ ama□zuz | 2 | 23.53% | 100% |
| ús□ik azzuga□ | 4 | 31.92% | 100% |
| amenzu | 7 | 24.56% | 90.68% |
| ús□ik nekwayes | 11 | 29.41% | 84.04% |
| ús□ik ezzuwa□ / zuwa□ |  |  | 46.43% |
| [tazuwa□t] | 8 | 27.95% |  |
| ka□ibi□ | 6 | 20.39% | 40.00% |
| ús□ik #1 | 18 | 23.97% | 47.06% |
| ús□ik #2 | 22 | 27.02% | 58.44% |
| tazuwa□t | 1 | / | / |
| taz□ubart | 1 | / | / |

For each named type, the local name of the dates is given, followed by the possible name of the palm [in brackets]. The identity is the average proportion of matching genotypes among accessions of a given named type and across the 17 nuclear microsatellites (Nuc Identity) and the chloroplastic minisatellite (Cp Identity).
\* one accession (3377) was identified as an identification error of the informer. When included, average identity across this set of accession is 79.41%.

### 3.1. Ethnographic and genetic analysis of date palms in Siwa region

#### 3.1.1. On the local named types of Siwa Oasis

The ethnobotanical field survey identified the existence of 18 named types in Siwa Oasis (Table 3). Based on ethnographic work, some were hypothesized as true-to-type cultivars, as they are supposed to only arise by planting offshoots, according to farmers’ accounts. Meanwhile, the fieldwork already allowed us to suppose that some of those named types are not true-to-type cultivars, as farmers recognized the possibility that they partially or totally arise from seeds.

To investigate this question, we calculated genetic identity among accessions of each given named type, based on 17 nuclear microsatellites and one chloroplastic minisatellite (Table S3). The named types □a□idi and alkak, the main cultivated and exported dates of Siwa, show proportion of identities of 97.6% and 100% at the nuclear level, respectively, and of 100% at the chloroplastic locus (Table 3, Figures 2-3). The named type alkak is locally known to come only from an offshoot, but different “qualities” can be distinguished, depending on growth conditions and age. The □a□idi is the emblematic “palm reproduced by offshoot”, which Isiwans always oppose to úšik, the seedlings. Although slightly different, the 13 □a□idi accessions cluster together (Figure 3), and we hypothesize that this subtle genetic variation is due to somatic mutations or genotyping error. Hence, the two Siwa elite named types are indeed true-to-type cultivars.

**Figure 2.**
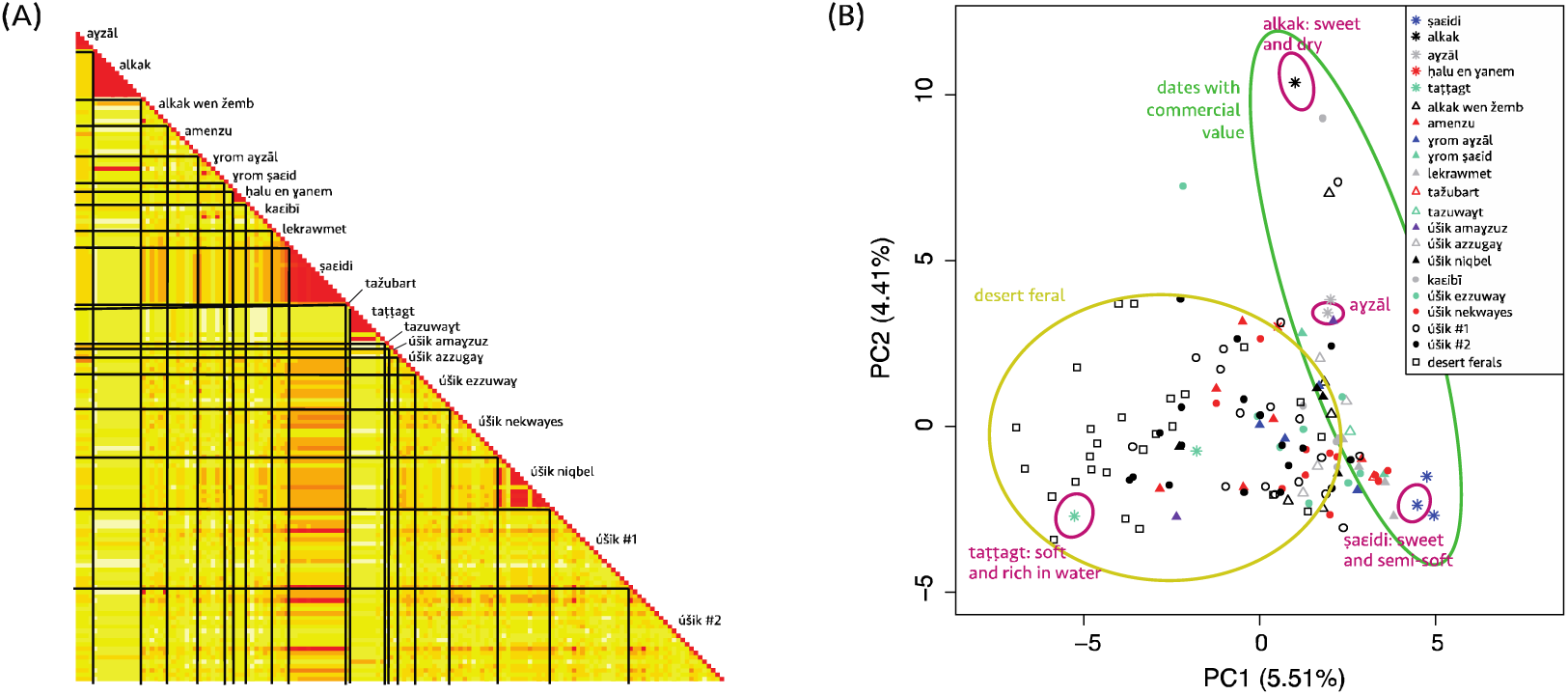
Intra-named type variation and structure in Siwa date palms. (A) Relatedness among the 149 cultivated date palms sampled in the current Siwa oasis, and genotyped across 17 nuclear microsatellite loci. Intra-named type variation is expected to be zero or near zero (red), in case of somatic mutations, while for ethnovarieties and local categories we expect a higher intra-named type variation (yellow) (B) Principal Component Analysis of 128 unique date palm genotypes from both the current oasis of Siwa and the abandoned oasis of the region. Variance explained by each principal component (PC) is provided within parentheses.

**Figure 3.**
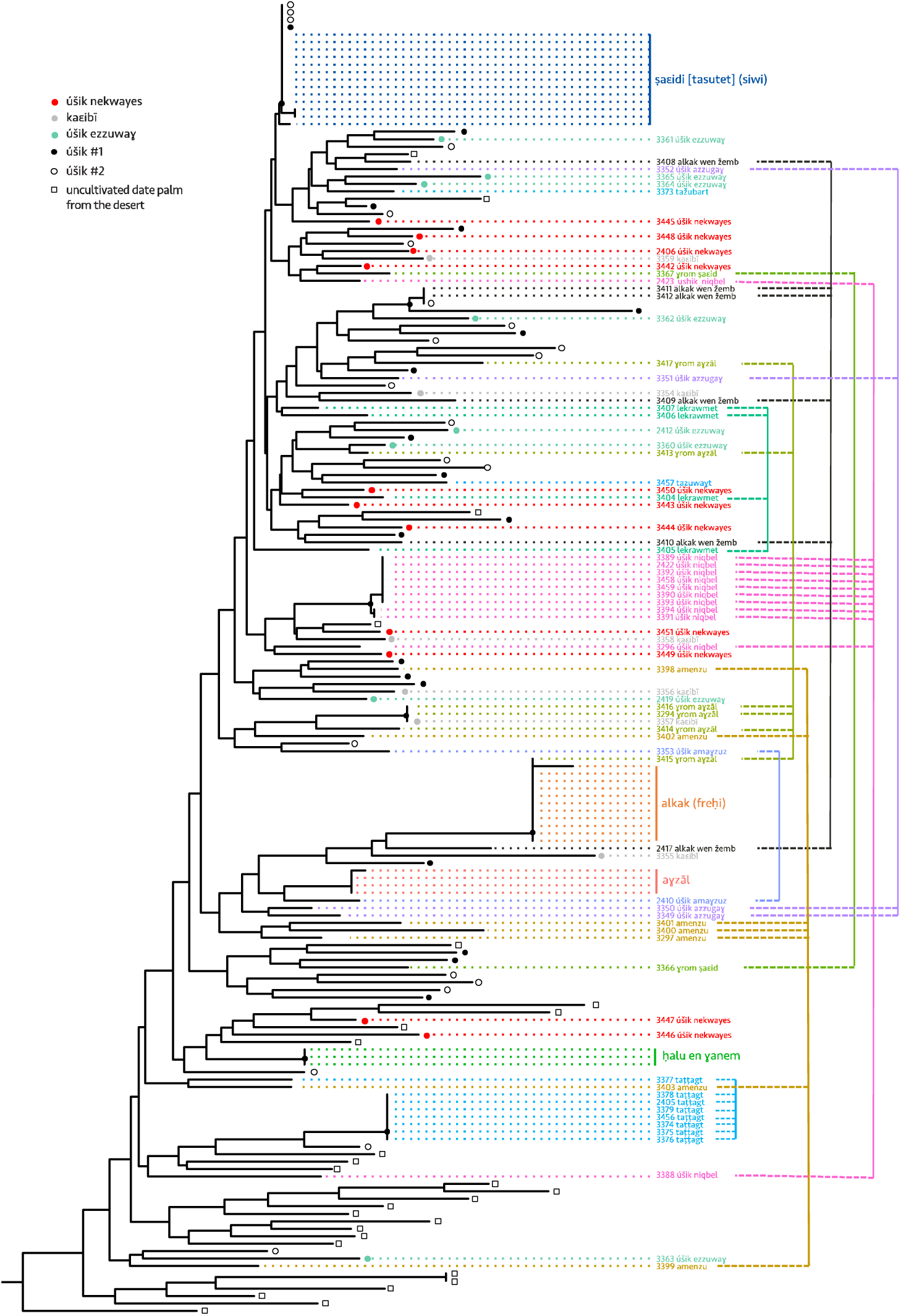
Genetic similarity of Siwa date palms based on Neighbor-Joining tree reconstructed from Nei’s genetic distances. On the right of the tree, colored vertical lines connect samples of the same named types. Nodes supported by bootstrap values >80% are indicated by black dots. The tree has been rooted with two *Phoenix reclinata* (not shown). The different samples of a true-to-type cultivar are expected to aggregate together, while samples of ethnovarieties would form different lines of clones, and samples of local categories would be scattered across the tree, denoting their lack of relatedness.

Have been also analyzed as true-to-type cultivar three other named types: a□zāl, ta□□agt, and □alu en □anem. The famous a□za̅l—rare but valued by the Isiwan as a tonic for lack of energy (Fakhry, 1990, p. 27) and for its aphrodisiac properties (for men)—is here represented by four accessions, three of which being identical and one showing a single allelic difference that we thus hypothesize to be a somatic mutation or genotyping error (Figure 3, Table 3, Table S2, Table S3). The named type ta□□agt refers to a very soft date rich in water, which name is taken from a maturation stage in the Siwa language: one half of the fruit is brown and ripe and the other half is yellow and immature. Because it keeps very poorly, it is always eaten on the spot, but is highly appreciated locally. It was studied using eight accessions, of which a single one (3377) appears to be very different from the other genetically uniform accessions (Figure 3, Table 3, Table S2, Table S3). The informant was a young farmer and we presume that he was wrong about the identification of this accession. Hence, we believe that ta agt is a true-to-type cultivar, even if different cc qualities of ta□□agt are reported by our ethnography. The fifth confirmed cultivar is □alu □anem (100% identity over the three studied accessions in both nuclear and chloroplastic loci, Figure 3, Table 3, Table S2, Table S3): not benefiting from much acknowledgement, relatively few farmers in Siwa know of its existence, but its long dates are highly appreciated by connoisseurs, so soft and sweet that the seed remains attached to the bunch when the fruit is plucked.

Further, we can confirm some named types as ethnovarieties: □rom a□za□1 and ús□ik niqbel indeed group some individuals having the same genotypes (clones) but also some that are isolated in different clades (Table 3, Figure 3). The first is said to resemble a□za□l, and the second □a□idi, but in both cases of lower quality.

An intermediate case is alkak wen žemb: the six samples are unrelated except two which are from the same clone, but coming from the same garden few meters away. Etymologically, alkak wen žemb is a “relegated,” “put aside”, i.e., a second class alkak, but it has always been, without much ambiguity, referred to as a cultivar and cannot come from a seed, according to farmers. Under the names lekrawmet, amenzu (etymologically “early” dates, (Laoust, 1932), or □rom □a□id, we only identified unrelated individuals (Figure 2, Figure 3, Table 3). These are clearly not true-to-type cultivars despite them being explicitly thought by local famers as never coming from a seedling; apparently, they often have been, although the occurrences may be too distant for local memory. They are thus interpreted as ethnovarieties.

Decisions are more difficult to reach in the following cases: ús□ik azzuga□, ús□ik ama□zuz, tažubart, and tazuwa t. The last two are only known to a handful of farmers, and we have only one sample of each, preventing an intra-type identity analysis. Tažubart is said to look like □a□idi, and tazuwa□t like ús□ik (zuwa□). The first two are more widely known, but two ama□zuz (etymologically “late” dates, (Laoust, 1932) samples do not allow to reach a conclusion. Regarding ús□ik azzuga□, we have two closely related palms from unrelated garden and farmers, but also two isolated samples. Local farmers say úšik azzuga□ are not (necessarily) reproduced by offshoot and can be from a seedling (úšik). They are sometimes described as “all úšik (seedlings) that yield red dates”, “azzuga” c meaning clearly “red” in *jlan en Isiwan*, the local Amazigh language. By crossing genetic information and ethnographic data, these four types thus all seem to be possible “local categories”.

We are quite certain that the following named types are neither cultivars nor ethnovarieties: obviously, and as expected, the inclusive local category of *úšik* in general (all female seedlings), but also úšik ezzuwa□ (*úšik* selected/cultivated with good reddish/dark dates), ka□ibī (*úšik* selected/cultivated with good dry dates), and úšik nekwayes (all *úšik* selected/cultivated with dates suitable for human consumption). Indeed, the field survey showed they refer to individuals sharing fruit characteristics and/or originate from seedlings, and the genetic analysis confirmed that samples are unrelated (Table 3; Figure 2-3). Four úšik nevertheless present the same microsatellite profiles as the Siwa widespread cultivar □a□idi (Figure 3) meaning that these were actually □a□idi but thought wrongly, by the sampler, to be accidental seedlings (because they were found in abandoned areas of the palm grove).

Following this identity analysis, we removed 58 genotypes from the initial Siwa dataset in order to only keep 128 unique genotypes of Siwa in downstream analyses.

#### 3.1.2. Structure of date palm diversity within Siwa region

To study the structure of the diversity of date palms in the region of Siwa, we studied named types and seedlings (úšik) from the oasis, but also uncultivated date palms from abandoned oases scattered across the desert surrounding Siwa (Figure 1). We found that the genetic diversity is mainly distributed between oasis samples *versus* samples from outside the oasis (i.e. the ancient abandoned oases in the desert), as the Principal Component (PC) 1 mostly draws apart those two types of date palms (Figure 2B). Identically, the NJ tree shows that the uncultivated date palms from the desert roughly form a distinct clade (Figure 3). Some accessions from the current oasis can nevertheless be found in the cluster formed by the desert uncultivated palms, mostly seedlings (úšik#1 and #2), but also named types, such as ta□□agt. The second PC opposes the two main cultivars, namely alkak and □a□idi. Alkak, and a few other c accessions, appear isolated from the other accessions of the region, including its relegated counterpart, alkak wen žemb (Figure 2B). The seedlings (úšik) do not form a distinct population. Here we differentiated the úšik from the gardens, that may be tended, depending on their use, irrigated, as other date palms of the garden, and pollinated (úšik #1) from those that are not (úšik #2). For farmers, there is no difference and based on the PCA, we could not separate them based on their microsatellite profiles.

### 3.2. Comparing the diversity of date palm in Siwa and worldwide

We compared 128 unique genotypes collected in Siwa Oasis and in the abandoned oases from the surrounding desert (Figure 1) with 219 date palms originating from North Africa and the Middle East (previsously published data: Moussouni et al., 2017; Zango et al., 2017; Zehdi-Azouzi et al., 2015), and nine newly genotyped *Phoenix theophrasti*. We investigated the population structure of these 358 unique genotypes with both Principal Component Analyses and Bayesian clustering, calculated F_ST_ between various populations and subpopulations, and estimated diversity in various populations and subpopulations.

#### 3.2.1. Partitioning and extent of the diversity in *P. dactylifera* and *P. theophrasti*

The Principal Components (PCs) 1 and 2 both separate *Phoenix theophrasti* from *Phoenix dactylifera* accessions (Figure 4A; Figure S2), accounting for the large differentiation observed between these two species (F_ST_ = 0.32, Table S4). Additional PCs do not provide notable results (Figure S3). On a side note, *Phoenix reclinata* being highly differentiated from other accessions (Figure S3), we excluded it from downstream analyses.

**Figure 4.**
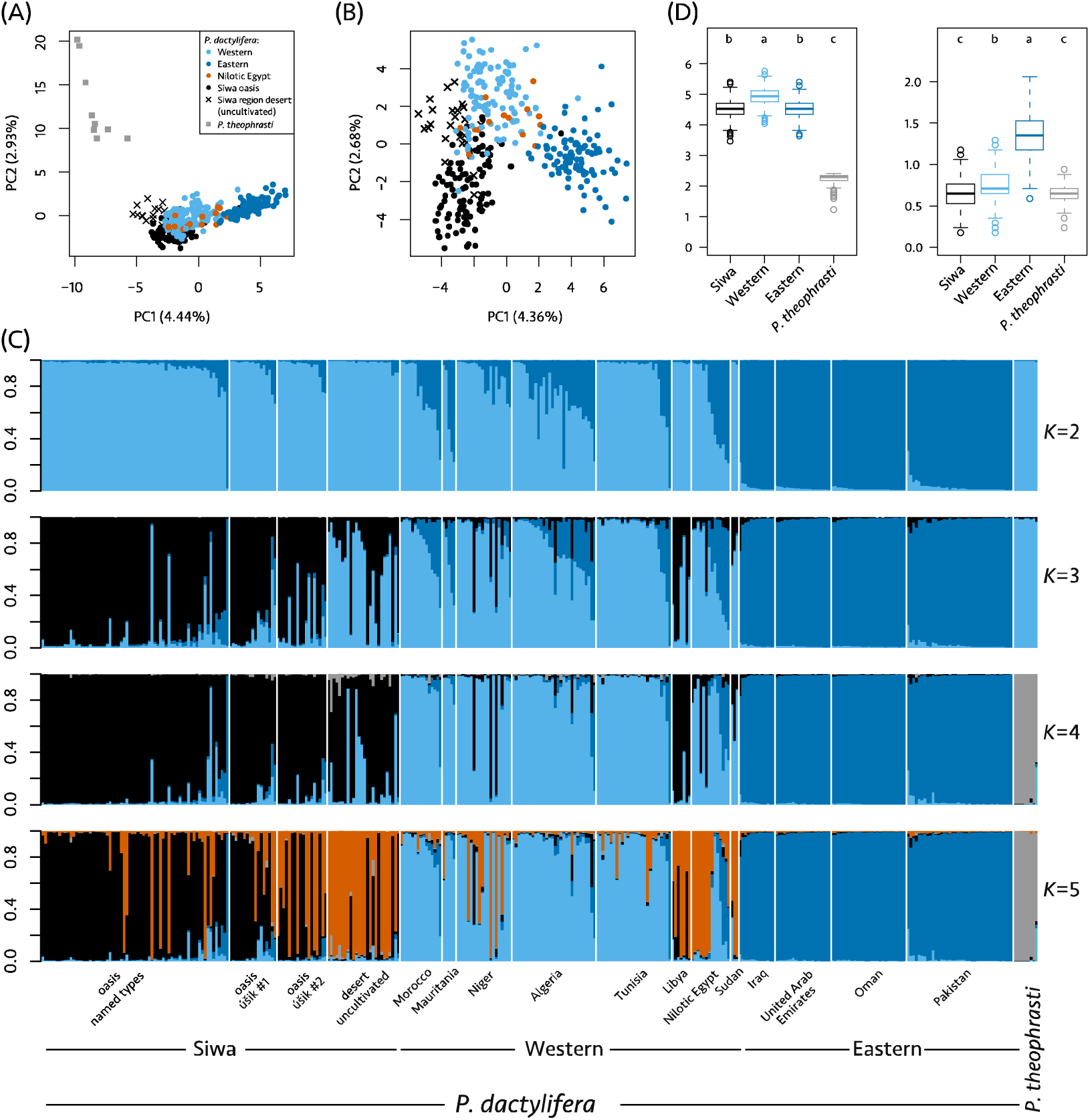
Worldwide structure of the diversity in date palms (*Phoenix dactylifera* L.) and *Phoenix theophrasti* Greuter. (A) Principal Component Analysis of 356 date palm and *P. theophrasti* accessions genotyped across 17 nuclear microsatellites (B) Principal Component Analysis of 347 date palm accessions genotyped across 17 nuclear microsatellites, (C) Admixture proportion in 356 date palm and *Phoenix theophrasti* accessions with K equal 2 to 5. Each individual is represented by a vertical bar partitioned into colored segments representing the assignment coefficient or ancestry, that is the estimated proportion of its genome derived from each cluster. (D) Allelic richness (left) and private allelic richness (right) calculated in four populations using the rarefaction method to equalize sample size across populations (haploid sample size = 18, bootstrap replicates = 1,000).

**Table 4.** Diversity estimates calculated in *Phoenix* theophrasti and various populations and sub-populations of *Phoenix dactylifera*.

| Populations | H<sub>O</sub> | H<sub>S</sub> | F<sub>IS</sub> (95% CI) |
| --- | --- | --- | --- |
| *P. dactylifera* | 0.56 | 0.63 | 0.10 (0.084 - 0.14) |
| Siwa | 0.56 | 0.60 | 0.060 (0.016 - 0.11) |
| Uncultivated desert | 0.55 | 0.64 | 0.14 (0.072 - 0.23) |
| Oasis | 0.56 | 0.57 | 0.015 (-0.028 - 0.066) |
| named type | 0.55 | 0.56 | 0.0077 (-0.036 - 0.054) |
| ús□ik #1 | 0.59 | 0.60 | 0.025 (-0.048 - 0.078) |
| ús□ik #2 | 0.57 | 0.59 | 0.02 (-0.060 - 0.12) |
| Eastern | 0.57 | 0.58 | 0.013 (-0.012 - 0.038) |
| Western (with Siwa) | 0.56 | 0.62 | 0.095 (0.064 - 0.14) |
| Western (without Siwa) | 0.56 | 0.61 | 0.081 (0.052 - 0.12) |
| *P. theophrasti* | 0.17 | 0.42 | 0.59 (0.26 - 0.83) |

H<sub>O</sub>, observed heterozygosity; H<sub>S</sub>, expected heterozygosity; F<sub>IS</sub>, inbreeding coefficient. Diversity estimates were calculated on western (or North African date palms) including or not Siwa accessions. Siwa population was split into two populations to oppose accessions sampled in the oasis (both named types and seedlings) with those sampled in abandoned oases in the surrounding desert. Further, accessions sampled in the oasis were further split into three populations: the named types and the seedlings collected in or nearby the gardens (ús□ik #1 and #2 respectively).

The Principal Component (PC) 1 describes the geography among date palm accessions, with Middle Eastern and North African accessions (including Siwa) being stretched from left to right (Figure 4AB). These two clusters are the first ones to emerge in the Bayesian clustering analysis (Figure S4, *K* = 2), and this partitioning is optimal according to the Evanno method (Figure S5). Eastern and western date palms are moderately differentiated, with F_ST_ = 0.088 or 0.084, whether Siwa accessions are included in the western cluster or not. The diversity among North African date palms is higher than among Middle Eastern date palms. Indeed, their levels of both allelic richness and private allelic richness, calculated with equal sample size, are significantly higher than that found in eastern date palms (Figure S6; Tukey’s tests, *P* < 0.05) and their expected heterozygosity reaches 0.62 while eastern populations display 0.58 (Table 4).

Interestingly, Siwa date palms overlap only slightly with the western and eastern clusters in the PCA (Figure 4AB). Similarly, in the Bayesian clustering analysis, Siwa accessions form a distinct cluster from *K* = 3 (Figure 4C). With an additional *K*, four clusters, corresponding mostly to *P. theophrasti*, eastern cultivars, western cultivars (excluding Siwa samples), and Siwa date palms, can be identified (Table S5). For the following sections, we thus consider those four populations, where Siwa and western populations are distinct. Siwa date palms appear more distinct from eastern (F_ST_ = 0.12, proportion of shared allele = 56.68%) than from western accessions (F_ST_ = 0.057, proportion of shared allele = 72.78%). In the Structure analysis they share only, on average, 2.73% of their ancestry with the eastern cluster, while 7.66% can be traced to the western cluster (Figure 4C, Table S5). The only Siwa accession with mostly eastern ancestry is an úšik ezzuwa□ (3363), collected in the heart of the old palm grove (in Jubba annēzi). Noteworthy, Siwa accessions are as diverse as eastern accessions in term of allelic richness (calculated using equal sample size, Figure 4D), and even more diverse in term of expected heterozygosity (Table 4).

On the one hand, the inferred ancestry constituting the Siwa cluster (Figure 4C, in black) is not only found in Siwa. Although it is almost absent in eastern cultivars (0.93% on average), it is substantial in western date palms (11.85% on average). It is prominent in the two Libyan accessions (Figure 4C): they share on average 89.86% of their ancestry with the Siwa cluster, the remainder being ancestry shared with the western cluster. On the other hand, we found that samples from Nilotic Egypt share ancestry with three clusters: mostly western (64.36%), but also eastern (18.36%) and Siwa clusters (16.81%). The evidence of shared ancestry with Siwa in the Nilotic Egypt cultivars is mostly driven by Hamra and Wardi cultivars (both from Aswan governorate), having both >70% of their ancestry shared with this cluster (a caravan road links Aswan to Siwa), while most other Nilotic Egyptian date palms share <1% ancestry with this cluster. The four Sudanese accessions share ancestry with both Siwa (28.30%) and western date palms (70.97%), and do not display ancestry traceable to the eastern cluster. We note that other North African countries share only a little of their ancestry with the Siwa cluster; Morocco (1.5%), Mauritania (1.26%), Algeria (3.56%), Tunisia (1.42%), the exception being Niger, with an average of 15.55%, driven by three cultivars (Kila72, Ja5, Jahaske2, all three from Goure, Zinder region, Southwest Niger, a new region for phoeniculture) with 94.80%, 92.30%, and 73.40% of ancestry attributed to the Siwa cluster, respectively.

#### 3.2.2. Shared ancestry between *P. dactylifera* and *P. theophrasti*

Structure results show shared ancestry between *Phoenix dactylifera* and *Phoenix theophrasti*. Indeed, eastern and western date palms share respectively 25.26% and 32.49% of alleles with the Cretan date palm. Structure results show shared ancestry between *P. theophrasti* and western date palms, but not with eastern date palms, except in Pakistan, probably due to recent introduction of African germplasm. Further, western date palms appear less distinct from *P. theophrasti* (F_ST_ = 0.32) than eastern do (F_ST_ = 0.40). While we reported above that western accessions display more private alleles than eastern ones (Figure S6), when we include *P. theophrasti* in the analysis, they display on the opposite fewer private alleles than eastern ones (Figure 4D). Indeed, over the 227 alleles identified across the 17 loci, 17 (7.5%) are shared by *P. theophrasti* and North African date palms, but not by Middle Eastern date palms. This shared ancestry may be evidence of either common descent or gene flows or both.

Curiously, Siwa accessions share even more ancestry with *P. theophrasti* than the other date palms (cultivars) of North Africa, including Nilotic Egyptian ones. Structure results indicate they have up to 29.3% of their genetic makeup shared with *P. theophrasti* cluster (on average: 1.24%, Table S5). They are also closer, and especially the uncultivated ones, to this wild Cretan relative on PC1 than western date palms are (Figure 4C, from *K* = 4), and share more alleles (35.58%) with this species than western do (32.49%).

Finally, Structure results show that there is also potential gene flow in the other direction, from *Phoenix dactylifera* to *P. theophrasti*. Indeed, two samples of the Cretan date palm display ancestry attributed to the date palm (Figure 4C).

#### 3.2.3. Partitioning and extent of the diversity within Siwa region

Siwa accessions can be further split in two distinct clusters, as seen previously (Figure 2B), and highlighted in both the Bayesian clustering (Figure S4, from *K* = 5) and the PCA (Figure 4AB). These two clusters mostly fit the pre-defined Siwa populations, with a cluster comprising mostly accessions from the current oasis (Figure S4, in black at *K* = 5), while the second comprises mostly uncultivated desert date palms from the abandoned oases (in orange at *K* = 5). Oasis and desert date palms are slightly differentiated (F_ST_ = 0.055). Uncultivated desert date palms are less differentiated from western date palms (F_ST_ = 0.051) than the cultivated Siwa date palms are (F_ST_ = 0.069). The accessions from the oasis, in contrast, are less differentiated from eastern date palms (Table S4).

Remarkably, the desert uncultivated accessions appear even less differentiated from *P. theophrasti* (F_ST_ = 0.28) than the accessions sampled in the oasis (F_ST_ = 0.35, Figure 4AC, Table S4). Structure results indicate an average shared ancestry of 4.10% between desert uncultivated accessions and *P. theophrasti*, while solely 0.33% of current oasis accessions ancestry can be traced to this species (Table S5, Structure at *K* = 4). Further, they share more alleles with *P. theophrasti* than Siwa Oasis cultivated do (38.89% and 34.45%, respectively). Those abandoned desert palms have almost no trace of the eastern ancestry (0.63%), contrarily to the named types from the oasis (3.07%). They are highly diverse in term of gene diversity (Table 4), and have, on average, more private alleles than the date palms from Siwa Oasis (Figure S6, Tukey’s test, *P* < 0.05).

The spontaneous seedlings found within and at the periphery of Siwa gardens (úšik #1 and #2, respectively) are found within the diversity of uncultivated and cultivated date palms of Siwa (Figure 4C, Figure 2B, Figure S6). They appear intermediate between named types and uncultivated desert palms in term of genetic makeup (Figure 4C, Table S5, Figure S4 at *K* = 5). Indeed, úšik #1 and #2 have 19.63% and 48.24% of ancestry shared with the Siwa desert uncultivated cluster, respectively (in orange at *K* = 5, Figure S4), and 12.40% and 74.66% shared with the named types, respectively.

#### 3.2.4. Diversity at the chloroplast minisatellite in Siwa and worldwide

We identified five alleles at the chloroplastic locus psbZ-trnfM (Table S2). Two of them were restricted to *Phoenix reclinata* and one of them to *Phoenix theophrasti* (Figure 4C). In *Phoenix dactylifera*, we identified the two previously reported alleles corresponding to the so-called occidental and oriental chlorotype (Gros-Balthazard et al., 2017; Zehdi-Azouzi et al., 2015). Eastern accessions mostly display the oriental chlorotype (94.90%, Figure S7). The occidental chlorotype is a predominant in the western accessions (71.74%), although there is a significant presence of the oriental one, reflecting seed-mediated gene flows from East to West more prominent than in the other direction (as shown by Gros-Balthazard et al., 2017; Zehdi-Azouzi et al., 2015). Most North African countries display a higher proportion of occidental chlorotype (Figure S8), except Mauritania as reported before (Gros-Balthazard et al., 2017).

Two accessions of *Phoenix theophrasti* display the date palm western chlorotype (Figure S8). These accessions also have ancestry that is mostly attributed to date palm in the Structure analysis (Figure 4C). This could indicate potential gene flow from the date palm to this wild relative, as previously reported (Flowers et al., 2019).

In Siwa, occidental and oriental chlorotypes are found in almost equal proportion (47.65% and 52.34%, respectively). Among the named types of the oasis though, the oriental chlorotype is slightly predominant (59.70%, Figure S8), while we found the opposite pattern in the Nilotic Egyptian cultivars (42.86%). As for the uncultivated date palms from the abandoned oases of Siwa desert region, we found that they display mostly the occidental chlorotype (74.1%).

## 4. Discussion

In this paper, we studied date palms from Siwa using a combined molecular population genetic and ethnographic approach, and in order to (1) better understand folk categorization in conjunction with local agrobiodiversity, and (2) infer the origins and the dynamic of the diversity found in this oasis and around, by comparing it to the worldwide date palm germplasm.

### 4.1. On the folk categorization in Siwa and its implication for surveying date palm agrobiodiversity

#### 4.1.1. Differentiating cultivar, ethnovariety, and local category

Our genetic analysis of intra-named type variability confirmed the existence of true-to-type cultivars as we found, for some named types such as the elite alkak, a 100% genetic identity across the 18 nuclear and chloroplastic loci. Nevertheless, we also pointed out the lack of genetic uniformity within other named types. Previous studies in date palms had already noted this dissimilarity and interpreted it as due to somatic or somaclonal mutations (Abou Gabal et al., 2006; Devanand and Chao, 2003; Elhoumaizi et al., 2006; Gurevich et al., 2005). Indeed, although offshoot propagation is alleged to be a true-to-type technique (Jain, 2012), somatic mutations can accumulate through generations of offshoots propagation. This has also been demonstrated in other crops, such as grape (Moncada et al., 2007) or cherry (Jarni et al., 2014). Further, when propagated through tissue culture, cultivars may accumulate somaclonal variation (El Hadrami et al., 2011; e.g. Medjool cultivar, Elhoumaizi et al., 2006). Here, we acknowledge that somatic mutations can indeed explain some variations. As a matter of fact, we interpreted slight genetic dissimilarity among □a□idi accessions as the result of somatic mutations, if not genotyping errors. Nevertheless, we also substantiate that the high genetic dissimilarity within some named types cannot solely reflect the existence of somatic mutations. Instead, it is the result of a cultivation practice that we proposed before, i.e. the incorporation of new clonal lines of seedlings under an existing name (Battesti & Gros-Balthazard et al., 2018).

Hence, using genetic data, we confirm the existence of distinct genetic statuses among the named types within an oasis territory included in this study: true-to-type cultivars, but also ethnovarieties, and local categories. Although we had previously hypothesized the existence of ethnovarieties (Battesti, 2013; Battesti, Gros-Balthazard et al., 2018), we here further substantiate this statement, using a larger number of named types and, for each, more samples. This study also highlights that a significant number of named types, beyond the very inclusive úšik and males, are neither true-to-type cultivars nor even ethnovarieties, but of the “local category” order.

Although we confirm the validity of these three notions, we also underline their limitations: they are hardly relevant for the farmers who perform these practices (see below *etic* vs. *emic*). Instead, the value of these notions lies in their ability to make the result of local agricultural practices intelligible to scientists.

This peculiar cultivation practice we observe in Siwa could also exist in other palm groves. Intravarietal variations have already been questioned for the bint aisha type sampled in different localities in Egypt (El-Assar et al., 2005, p. 606), for other named types in Libya (Racchi et al., 2013), and for samples from different countries or different regions (Khanamm et al., 2012, p. 1240). Authors have also noted the apparent dissimilarity between individuals of the same variety in the same oasis most often attributing it to somatic mutations (or, in morphological characterization studies, to cultivation conditions, care and age of the date palm, for instance in Rhouma, 1994). Here, the case is different in that the intravarietal variations are internal to an oasis, and they are the result of a voluntary phoenicultural technique (Battesti, 2013). Such a farming practice producing ethnovarieties could also explain the “unknown mechanism” behind the fact that Medjool/majhūl, the famous Moroccan variety, is not a “genetically uniform” clone (Elhoumaizi et al., 2006, p. 403).

#### 4.1.2. Determining the number of named types

Surveying the precise number of named types that refer to distinct local date palms in an oasis is an already complex operation, as in Siwa (Battesti 2013) where given lists of named types can refer to varying degrees of inclusiveness (in no hierarchical order of exclusive taxa, unlike what Brent Berlin’s ethnobiological theory of taxonomic categories implies: Berlin *et al*., 1973, 1974). However, in a system with cultivation methods based on massive vegetative reproduction, the identification of the accurate number of named types should have been sufficient to correctly estimate the number of genotypes. Until now, therefore, and at best, the overestimation of the agrobiodiversity of the date palm at the regional level had been considered, taking into account the phenomenon of synonymy (a cultivar takes another name by changing oasis), somewhat offset by homonymy (the same name is used in different oases to designate a different cultivar) (Battesti, 2013; Battesti et al., 2018). In a single oasis, Siwa, one of the difficulties of the field survey resides in massive local synonymy, including for plant names and in particular for date palm varieties, as already mentioned (Battesti, 2013; Battesti et al., 2018). The first hypothesis addressing such a synonymy is that a landlocked Berber-speaking community should promote its export products by adding Arabic trade names. Another explanation not to be underestimated is the co-presence on the same territory of Arabic-speaking minorities (sedentary Bedouins, especially Awlad ‘Alī, Battesti & Gros-Balthazard et al., 2018) who use these Arabic names; for example, rather than the names tasutet/ alkak/ úšik they will systematically use (or even only know) the names □a/□idi/fre□ī/ azzawī to denote the same palms. Besides, we observed that local practices of categorization and integration of seedlings can lead, as explained above, to a massive phenomenon of underestimation of agrobiodiversity for an uninitiated external observer and all assessments have stumbled over this obstacle (even the very recent *Atlas of date palm in Egypt*, El-Sharabasy and Rizk, 2019).

In Siwa, our ethnobotanical survey revealed the existence of 15 to 20 named types (Battesti, 2013; Battesti & Gros-Balthazard et al., 2018). The exact number is difficult to pinpoint as the field survey reveals that farmers offer local, more or less shared, quality distinctions, even for dates that we depicted as true-to-type cultivars (alkak and ta□□agt for example). Therein, a higher alkak is called “alkak n amles”, meaning smooth or wrinkle-free alkak dates, and a lower alkak with smaller dates, and three times cheaper than the upper one, is called “alkak nifu□en”.

Fifteen to 20 named types in Siwa is not a lot compared to the number of named types described in other oases. Assessing the agrobiodiversity of date palm is a difficult exercise, and carried out using non-comparable competing methodologies (Jaradat, 2016). As a result, it is difficult to establish the terms of comparison of agrobiodiversity between oases. We can provide some comparative data: for instance, in the Jerid region of Tunisia (about twice the area of old palm groves compared to Siwa), there is a collective collection of more than 220 varieties (Battesti, 2015; Rhouma, 2005, 1994). (Some of them may very well be ethnovarieties or local categories, but this hypothesis still has to be checked *in situ*.) Smaller oases have about the equivalent number of named types: 18 in Sokna (al-Jufra, Libya) for a cultivated area half as small as Siwa (Racchi et al., 2014), 18 in Kidal (Northern Mali) but for only 4,000 date palms (Babahani et al., 2012), 22 in el-Guettar (Tunisia) for only 3,000 date palms (Ben Salah, 2012). Hence, we could believe that the agrobiodiversity is relatively low in Siwa, if considering a conventional system where named types are thought to be genotypes. Nevertheless, with this combined ethnobotanic/genetic approach, we showed that this assumption is wrong, and that, despite a low number of named types compared to other oases, the number of genotypes in Siwa is high and so is the overall diversity.

#### 4.1.3. Etic vs. emic categorizations

Two main difficulties arise when assessing local date palm agrobiodiversity. On the one hand, date palm is supposedly reproduced by local farming communities through vegetative propagation (offshoots), but we demonstrated that apparently there are a number of exceptions in Siwa (all the non-true-to-type cultivars) that are not formally documented at the local level. On the other hand, and this explains the first point, the local ways of conceiving this living material and its qualities, of categorizing it (by form) do not quite match with biologists’ ways of conceiving it and categorizing it (by genes) (Battesti, 2013). This difficulty is a classic conflict for anthropologists, also known as a disparity of *etic* vs. *emic* categorizations (Olivier de Sardan, 1998). Deeply characteristic of human societies, categorization processes are also at the heart of both mundane and scientific thoughts and practices. Naturally, our aim is not to use our paradigm (genetics) to try to evaluate the paradigm of indigenous local knowledge (Roué and Nakashima, 2018), but to translate the latter: for social sciences, there is no such thing as one science but several incommensurable sciences, and multiple modes of existence coexist responding to various forms of veridiction (Latour, 2013). In the *emic* version (the local point of view), the differentiation between cultivars/ethnovarieties (which farmers assimilate) and local categories is clearly thought out. However, it is not so clearly expressed: farmers can list all named types at the same level even if they do not refer to the same object classes. It is worth bearing in mind that for local farmers, reproducing by offshoot is the rule. Naming/identifying a seedling date palm after a known cultivar (ethnovariety process) is a possible (and appropriate) practice but of an exceptional occurrence on a human life scale. Hence the difficulty for local farmers to know if, for instance, all the úšik niqbel are cultivars (which they tend to present as such) or ethnovarieties. Although they do not remember seeing an úšik n gubel coming from a seed, this has apparently been the case, perhaps several decades or generations ago (a date palm easily outlives a human being). The question is of little relevance to them, since the palm trees in question behave and have/produce the same form. But, the other process, categorizing a date palm from a seedling, with reddish/dark dates for instance, as having a valuable production and naming/qualifying it, as úšik ezzuwa□ for instance, is a more common experience (local category process). Hence, for local categories, we have to reverse the point of view. For example, some date palms do not bear reddish/dark dates because they belong to the variety úšik ezzuwa□ or zuwa□, but the fact that their dates are reddish/dark qualifies them as úšik ezzuwa□ or zuwa□. As scientists, embedded in our *etic* point of view, the difference between ethnovarieties and local categories may seem thin as both result in genetically unrelated individuals (with a few clones for the former if sampling is sufficient), hence the necessity to consider cultivation practices and local knowledge. Indeed, to distinguish between what can be objectified as an ethnovariety or a local category in our samples is not easy. A clear choice for cultivar true-to-type can be made. Another clear choice is possible for an ethnovariety in the case of x clones plus one or two outsiders (possibly also clones, we did not face the case). When all individuals are different from each other, there is a greater chance of having a local category (to be confirmed by the way farmers talk about it), especially when there is a minimum local consensus. Assessing this consensus is obviously difficult, since no one in the community will use our *etic* way of thinking the diversity by cultivar/ethnovariety/local category. Informants just state it as “this date palm, it is an xxxx” [a named type]. Even more complex is the case of the rare ethnovariety, which is little known, but real, so few individuals are sampled that there may not be any clones in the sample.

It is therefore preferable to consider the distance between the ethnovariety and the local category as a continuum (Figure 5). Indeed, among the 18 named types we analyzed here, we found a gradient of intracultivar relatedness: some named types corresponded actually to unique genotypes while some did not, as evidenced in our previous study (Battesti et al., 2018). It is worth noting that an expanded sampling could show that what we believe today is a true-to-type cultivar is in fact an ethnovariety, by discovering a new line of clone under the same name. Further, it could allow to identify lines of clones for the inconclusive alkak wen žemb, lekrawmet, amenzu or □rom □a□id to confirm them as ethnovarieties. To formally confirm that a given name is either a true-to-type cultivar, an ethnovariety or a local category, it would require that the 200,000 to 250,000 date palms of Siwa to be all genotyped. This is obviously not conceivable.

**Figure 5.**
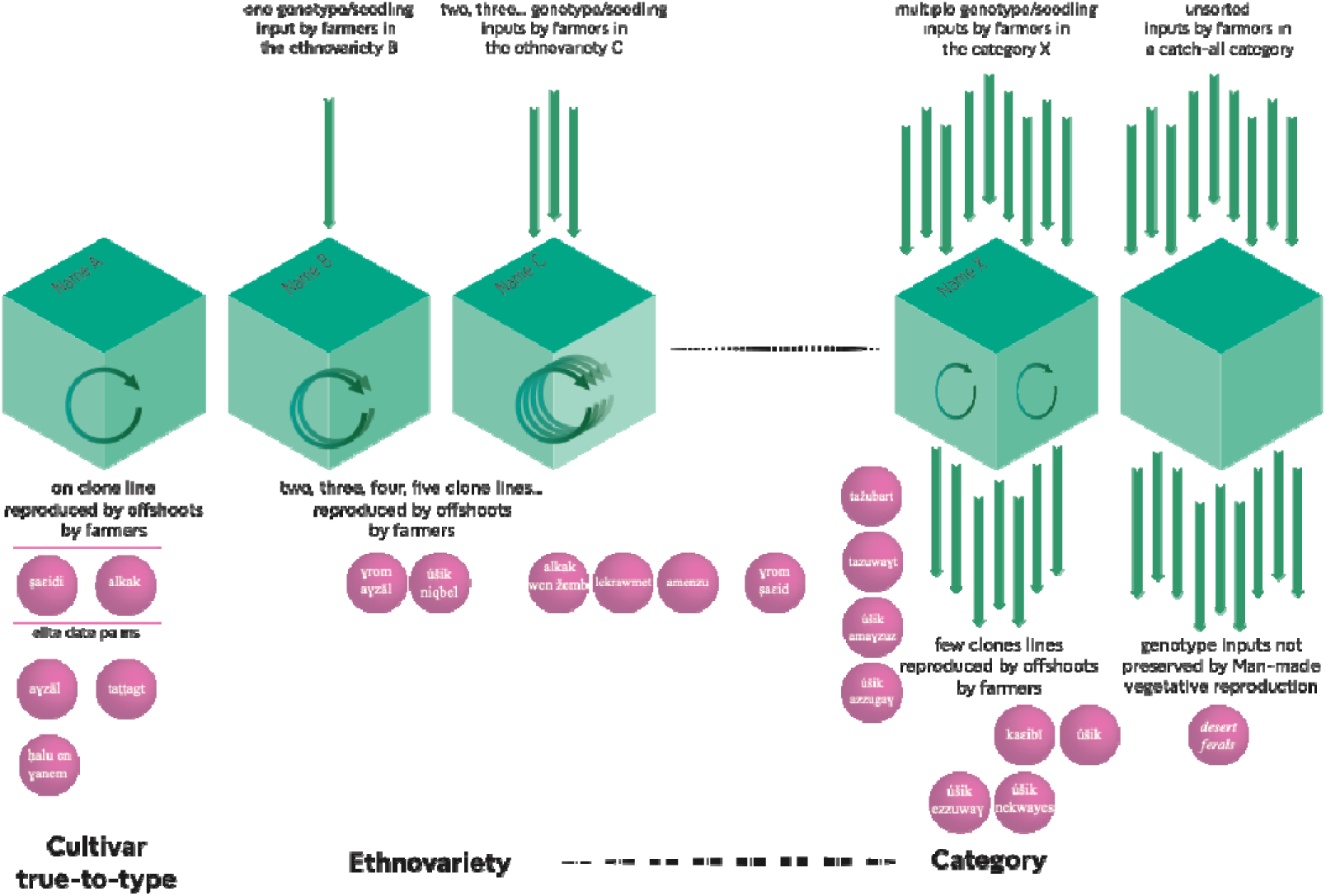
Converging local categorization of date palms in Siwa (*emic*) and genetic categorizations (*etic*) with the distribution of the named types identified by Isiwan. For genetic analyzes, there is a continuum between ethnovariety and local category, while for local farmers, the rationale is quite different (genotype vs. phenotype approaches). For farmers, ethnovarieties imply the idea of identity, when local categories refer to the idea of a qualification (to qualify a date-palm of…). Nevertheless, cultivars, ethnovarieties, and local categories are listed by Isiwan as if they were equivalent.

### 4.2. Usefulness of the folk categorization

What are the explanations for such a complex local system for categorizing date palms in Siwa? They might be both cognitive and agricultural. First, as with all classification, the aim is to recognize, to stock a huge amount of information, and to be able to mobilize it easily in order to guide action and communicate (knowledge, experience, etc.). This system must therefore be shared. It is a hypothesis supported, for example, by Meilleur in the context of ethnosciences: “such classifications are functionally linked to the effective storage, retrieval and communication of large quantities of information relating to the animal and plant worlds” (Meilleur, 1987, p. 9-10). The utilitarian nature of folk biological classifications had already been discussed by Eugene Hunn (1982). Date palm cultivation in Siwa is largely dominated by a few “elite” types (probably for centuries, an integration into the Saharan trading network, Battesti, 2018, 2013). Here we found that they are apparently true-to-type cultivars, despite their prevalence and therefore the mechanical possibility of becoming an ethnovariety. The ethnovariety and local category system makes it possible to “put in order” the profusion of other date palms, less commercially valued, while not multiplying the denominations for the same characteristics.

Secondly, a peculiar system allows for a peculiar action on the world, in this case a fairly flexible management of agrobiodiversity. McKey *et al*. (2010) consider that cultivation of clonally propagated plants represents a singular system that has to date been largely neglected, and that a key component of strategies for preserving the adaptive potential of clonal crops is the maintenance of mixed clonal/sexual systems. Their model is the cassava (*Manihot esculenta* Crantz), a eudicot grown as an annual plant in tropical and subtropical regions. To date, cassava is the only clonal crop for which in-depth information exists on how mixed clonal/sexual systems work (*ibid.*). We propose here another crop model, quite different as the date palm is a perennial plant that lives decades or even over a century (Chao and Krueger, 2007). This critically impacts farming practices, especially those inherent to propagation (by seed or offshoot, sexual or clonal). However, it is very likely that maintenance of a mixed clonal/sexual system is a local key strategy for managing and preserving an agrobiodiversity of date palms in Siwa. And this is made possible for Siwa farmers because they developed a classification system that enables them to do so.

How does it usually work—or how do we think it works? Usually, North African of Middle Eastern oasis farmers have important stock of cultivars (true-to-type) reproduced vegetatively, and a possible gene pool of seedlings, with their own possible use (fodder, handicraft, etc.)—named depending on places *khalt*, *sheken, degla* (Battesti, 2005), *rtob* (Ben Salah, 2012), *saïr* (Peyron, 2000), *sayer* (Naseef, 1995), *qush* (Popenoe, 1913), *dabino* (Zango et al., 2016), etc., or *úšik* in Siwa. From this stock is sometimes drawn on occasion a new seedling then socially recognized by the granting of a new name and then vegetatively reproduced: it enhances the initial collection.

We have demonstrated that in Siwa, the local categorization system including ethnovarieties and local categories—part of the palm domestication at ethnographic scale—allows an even more flexible management of agrobiodiversity and gene flow to the cultivated pool, as any new seedling integrated into the procession of cultivated date palms is likely to be classified within a pre-existing named type, although not sharing the same genetic heritage in common (genotype), but sharing, from a local point of view, the same form or the same characteristics (phenotype).

### 4.3. A unique and high genetic diversity in Siwa date palms

Siwa date palms form a partially distinct cluster from western accessions, rather than a subset of the diversity found in North Africa as it might have been expected. Some alleles are found uniquely in the region. The level of allelic richness in Siwa (a tiny region) is strikingly as high as that found in all eastern accessions, originating from a large geographic area from the Near East through the Arabian Peninsula and as far as Pakistan. We note that Libyan cultivars also belong to what we can call a third genetic cluster of modern date palm germplasm. This uniqueness of date palms from the Libyan Desert has never been reported before. In fact, very few studies have compared accessions from Siwa with other date palms. Gros-Balthazard *et al*. (2017) mentioned that the cultivar “siwi” from Siwa (here called □a□idi) did not cluster with the Middle Eastern accessions nor with the North African accessions, but it was interpreted as a hybrid origin. Identically, it does not cluster with other Egyptian accessions in a study based on AFLP (El-Assar et al., 2005). A seed morphometrics analysis shows that it clusters with date palms of mixed origins (Terral et al., 2012). Nevertheless, both a limited sampling in Siwa and an incomprehension/ ignorance of the cultivation practice had hampered those previous researches to pinpoint the singularity and high diversity of Siwa date palms.

Three non-mutually excluding hypotheses could explain the high genetic diversity observed in Siwa date palms. First, a highly diverse but yet unidentified source of diversity may have contributed to date palms in the Libyan Desert, thus making them both highly diverse and unique, compared to other regions (see section 4.4). Second, the cultivation practices that we here describe may have maintained/promoted this high diversity. Indeed, a sole clonal propagation leads to a loss of diversity (McKey et al., 2010). In Siwa, we found a complex system where what we thought were cultivars (clones) are in fact ethnovarieties or local categories, which in turn account for a higher diversity than expected at first sight. It is thus possible that in Siwa, more than in other oases, sexual reproduction being more common, there is a more limited loss of diversity through time. This hypothesis remains to be tested by ethnographic survey in other oases in order to test whether sexual reproduction is more prevalent in Siwa than elsewhere. This leads us to the third hypothesis: the high diversity found in Siwa could reflect, rather than a reality, a sampling bias. Indeed, our sampling, considering practices and knowledge of farmers, may have led us to sample an appropriate representation of the existing diversity of Siwa. On the opposite, the non-Siwa date palms included here have been sampled without such an ethnobotanical survey, and may in turn only be a poor representation of the actual diversity. Hence, if such a sampling methodology were applied everywhere, we may very well discover more diversity elsewhere too.

We identified that date palms in the current oasis of Siwa and in the ancient, abandoned palm groves in the surrounding desert constitute two subpopulations. While we identified potential gene flow from eastern accessions at both chloroplastic and nuclear level in the date palms from the current oasis, accessions from the abandoned palm groves may have received less diversity from this population. Additionally, uncultivated palms from the desert share more ancestry with *P. theophrasti* than both western and Siwa Oasis date palms. Locally, these desert date palms are considered as úšik but are also specifically called igizzã (sing. agzzu). Our ethnographic survey revealed that they do not seem to be used by the farmers as a reservoir of diversity. Some of these abandoned oases could nevertheless constitute casual date harvest sites, and hence seeds from the harvested fruits could potentially end up as úšik in the cultivated palm grove. This could explain the intermediate genetic profiles of úšik #1 and #2: their diversity seems in between that found in the current oasis and the abandoned oases.

Siwa date palm origins are unknown, but their presence in Siwa dates back at least to the 5^th^ century (mentioned by Hellanicus of Mytilene, see above); the oasis being then an independent state related to the Libyan world, well established as an essential trade and religious hub with Libyan, Egyptian and Hellenistic influences (Kuhlmann, 1999). Declining or abandoned at the end of the first millennium, the oasis was probably recolonized in the 11^th^ or 12^th^ century (*a priori* by Amazighs from Libya, then Arabs, see Battesti, 2013), but we do not know if this was done by conserving the original stock of date palm trees or by introducing new plants/cultivars, or both. In the present study, we found that the date palm population of Siwa is unique. The opportunities for a melting pot of genetic richness of date palm were there from the (known) “origins” of Siwa, because since the 6^th^ century BCE, the realm of “two deserts” (eastern and western deserts, from a Siwa-centered point of view) covered the eastern Saharan oases as far as Bahriyya, but probably also the oases along the tracks leading to Nubia, and in the other direction of course Awjila, but also probably to the south at Kufra Oasis (Kuhlmann, 2013). Nevertheless, the connections are not enough: we would have to estimate on the one hand what the dispersion flows of the domestic date palm in these oases have been, and on the other hand whether we are dealing with an entry of genes into Siwa that is involuntary—by dispersed seeds from elsewhere—or voluntary—and there probably by the contribution of off-shoots. For the latter hypothesis, we must consider a model of connectivity from one place to another, from oasis to oasis, the “stepping stones” model of the landscape ecology (Burel & Baudry, 1999), a “functional connectivity” (Battesti, 2018). The genetic makeup in Siwa, like that of other North African populations, may originate from a mix of Middle Eastern date palms and *P. theophrasti*; an unknown ancestral gene pool may also be involved (see below).

### 4.4. Revising the history of date palms origins and diffusion

Using 17 nuclear microsatellites and one chloroplastic minisatellite, we detected the previously described differentiation between date palms from North Africa (the so-called western population) and those from the Middle East and Pakistan (the so-called eastern population). Our results also corroborate the existence of gene flows between these two clusters, mostly from East to West (Gros-Balthazard et al., 2017; Hazzouri & Flowers et al., 2015; Zehdi-Azouzi et al., 2015).

#### 4.4.1. The contribution of *P. theophrasti* to North African date palms: insights from Siwa date palms

Date palms were domesticated at least in the Persian Gulf region, followed by diffusion to North Africa (for review, Gros-Balthazard et al., 2018). A recent study, based on whole-genome re-sequencing data, indicated that North African date palms have been introgressed by the Aegean *Phoenix theophrasti*, and today, about 5-18% of their genome originates from this wild relative or a *P. theophrasti*-like population (Flowers et al., 2019). Here, we also identified potential evidence of this introgression. Indeed, the Bayesian clustering identified shared alleles between these two populations, and the differentiation between western date palms and *P. theophrasti* is reduced compared to that between eastern date palm and this wild relative. We however note that none of the 121 North African and 156 Siwa date palms display the *P. theophrasti* unique chlorotype (five repeats of the dodecanucleotide minisatellite, Pintaud et al., 2013). This corroborates previous results showing that plastid and mitochondrial haplotypes characterizing *P. theophrasti* are absent in the 25 sequenced North African date palms (Flowers et al., 2019). This has been interpreted as an asymmetry in the direction of the interspecific cross: gene flows from *P. theophrasti* were pollen-mediated (*ibid.*). We note nonetheless that the possibility of a plastome-genome incompatibility (Greiner et al., 2015) cannot be ruled out.

Although a major event in the diffusion and diversification of cultivated date palms to North Africa, this introgressive event remains puzzling, especially in terms of localization (Flowers et al., 2019). Indeed, today, *P. theophrasti* is not distributed in North Africa, but is found in the Aegean region, mostly in Crete (Barrow, 1998), but also in coastal Turkey (for review, Boydak, 2019). One hypothesis explaining the introgression by *P. theophrasti* is that it had a much wider distribution in the past, encompassing North Africa and/or the Levantine region (Flowers et al., 2019). In the literature, authors consider it is a tertiary relict species (Vardareli et al., 2019). Here, we indeed found a high inbreeding coefficient that could reflect a severe population bottleneck due to habitat loss. Nevertheless, we lack positive evidence for the presence of the Cretan date palm in the region and this hypothesis remains unconfirmed.

In this study, we found that date palms from the region of Siwa share on average more alleles with *P. theophrasti* than other western varieties. This could indicate that North African date palms were introgressed by *P. theophrasti* or a *P. theophrasti*-like population in this region. As a matter of fact, there were tight historical links between Crete and the region, and although this is only speculative, and we do not have the means to propose a chronology, historical elements may suggest avenues for reflection and hypotheses. For instance, the Minoan culture, that had developed in Crete, interacted with the mainland Eastern Mediterranean, including New Kingdom Egypt, most intensively in the second half of the second millennium BCE (but evidence for earlier contacts exists). Indeed, there is extensive archaeological and textual evidence for trade and diplomatic relationships, as well as the presence of Minoan artisans in Egypt, for instance in Avaris/Tell el Dab’a in the eastern Nile Delta (Bietak, 2005; Bietak et al., 2007). Minoan Crete exported timber, textiles, probably olive oil and other products. These exchanges seem to have lasted a few centuries, during which time Cretan date palm products may have been brought to Egypt (Bietak, 2005). Half a millennium later, Siwa became the center of a possible realm of “two deserts” (Kuhlmann, 2013), which center included the now abandoned oases in which we sampled and others further west from Shiyata (Šiyata) via Jaghbūb to Awjila/Jīlō in Libya. At least from the 5^th^ century BCE onwards, Siwa was one of ō the large hubs, markets and centers that controlled a major part of all movements coming from and going in their directions (Rieger, 2017, p. 52), while going unmentioned in sources from the Nile valley (Kuhlmann, 2013). Siwa was caught between a marked autonomy resulting from its singular Libyan-Berber identity and the inevitable influence exerted on it by the important surrounding political realities: Pharaonic and then Graeco-Roman Egypt on the one hand, and the Greek colonies of Cyrenaica on the other (Struffolino, 2012). The contacts were close in particular with the Western Pentapolis (the eastern part of Cyrenaica), and the most important colony, Cyrene. Greek inscriptions by Greek and even Cretan workmen during the late fourth to early third century BCE were found in Siwa (Aldumairy, 2005; Kuhlmann, 2013). In short, there was an Egyptian cult with a deity venerated in the temple taken from the Nile Valley in a Libyan oasis whose history was shaped by the Greeks (Bruhn, 2010). Did one of the many Athenians (Colin, 1997) or Greek delegations in general who presented their offerings to the patron god of Siwa bring Cretan palm pollen? Or were they just merchants, as Kuhlmann (2013) argued that trade, not oracles, were behind the oasis’ appearance on the historical scene?

We note that date palms from the abandoned oases of Siwa show an even closer affinity to *P. theophrasti* than the palms from the current oasis. Again, we can only speculate about the processes resulting in this pattern. One hypothesis is that those palm groves were abandoned shortly after introgression by the Cretan date palm, and their genetic makeup may have evolved independently from other date palm populations. In addition, date palms in the current oasis could have had their *P. theophrasti* ancestry diluted by gene flows from other date palm populations, especially the eastern ones.

#### 4.4.2. The Siwa region at the diffusion crossroads of the domestic date palm?

Siwa was in early contact with the Greek colonies of the Cyrenaica, but also with Egypt and the Libyan Sahara. For the diffusion of the date palm, its position must have been essential.

In Egypt, unequivocal evidence for the economic and cultural importance of dates and for the local cultivation of the date palm for its fruits begins with the New Kingdom in the mid 2^nd^ millennium BCE (Tengberg, Newton 2016). However, there is now evidence for the presence of date palm products in Egypt, at least in contexts related to the crews working for the building of royal pyramids during the Old Kingdom (ca. 2700-2100 BCE); administrative texts document that two regions (nomes) located on the Mediterranean coast provided dates to workers under the reign of Khufu (around 2600 BCE, Tallet, 2017), while a very small amount of date palm remains were found at Giza (2700-2100 BCE) (Malleson, 2016; Malleson and Miracle, 2018), and at the site where the texts were found (Wadi al-Jarf, a Red Sea harbor site, Newton, unpublished data). Whether the dates were produced locally, in the Nile delta, or imported through potential Mediterranean harbors from somewhere in the Eastern Mediterranean remains unknown.

The extension of the route leading west from Siwa, via Jaghbūb, Awjila, to Fezzan, and plausibly onwards appears to date to the late second or early first millennium BCE (Mattingly, 2017, p. 8). In the Fezzan (Libya), the Garamantes developed into a Central Saharan state (*ibid*. p.15) and were involved in separate bilateral trading arrangements with neighboring states and peoples to the north and south (*ibid*. p.19). Late pastoral period sites in the Fezzan do suggest some contact with agriculturalists: date stones were recovered dating to the end of the second millennium BCE, but could have been “exotic” imports from Egypt. From the beginning of the first millennium BCE, the occupants of Zinkekra were cultivating a range of crops characteristic of Near Eastern, Mediterranean and Egyptian farmers, based on emmer wheat, barley, grape, fig and date (Pelling, 2005, p. 401). These are some of the earliest evidence for oasis agriculture in North Africa (Van der Veen and Westley, 2010). The realm of the “two deserts”, in Siwa, could have been in the first millennium an important node in the date palm exchange network between Egypt and Libya.

#### 4.4.3. A missing contributor to the modern genetic makeup

North African date palms display a higher diversity than Middle Eastern ones (Gros-Balthazard et al., 2017; Hazzouri & Flowers et al., 2015). Flowers *et al*. (2019) showed that the more genomic regions are introgressed by *P. theophrasti* or a *P. theophrasti*-like population, the higher their diversity, concluding that the excess of diversity found in North African date palms can be explained by the introgressive event(s) by this wild relative. Our results corroborate these findings, as we found that the fraction of alleles uniquely found in North Africa is reduced when *P. theophrasti* is included in the analysis.

Nevertheless, both Flowers et al. (2019) and our data indicate that there could still be an unknown population that contributed to the modern date palm germplasm. Indeed, in genomic regions with no or limited *P. theophrasti* introgression, the diversity in North African date palms remains higher than that in Middle Eastern date palms (Flowers et al., 2019). Further, of the two main chlorotypes described in date palms (Pintaud et al., 2013), we ignore the origins of the so-called occidental one, prevalent in western date palms. Indeed, the few relictual wild date palm populations bear the so-called oriental chlorotype (Gros-Balthazard et al., 2017), and *P. theophrasti* an even more divergent type (Pintaud et al., 2013). Here, we could also identify diversity that is unique to the North African date palms, especially in Siwa. The origins of this unique genetic makeup are unknown. It could, like other North African date palms, derive from a mix between Middle Eastern date palms and *P. theophrasti*. Indeed, demographic events, such as population bottlenecks, or selection can lead to drastic changes in allelic frequencies, while mutations may introduce new alleles. Another hypothesis, reinforced by results obtained at the chloroplastic loci, is a contribution from not only Middle Eastern date palms and *P. theophrasti*, but also from a yet identified North African *Phoenix* populations.

So far, there is no evidence of date palms or other *Phoenix* populations in North Africa before the establishment of oasis agriculture. Pre-domestication *Phoenix* remains attributed to *P. dactylifera* have only been found in the Levant and in Iraq (Henry et al., 2011; Liphschitz and Nadel, 1997; Solecki and Leroi-gourhan, 1961). In contemporary times, wild date palm populations are exclusively known in the mountainous regions of Oman (Gros-Balthazard et al., 2017). The natural distribution of *P. dactylifera* (before its domestication and diffusion) is unknown (Barrow, 1998). It likely covered at least the Middle East, and may have shrunk, in connection to climate change (Collins et al., 2017). In Egypt, it is possible that the date palm was growing wild, or that it was grown for ornamental purposes (more precisely, not mainly for its fruit), perhaps from the 4^th^ until the mid 2^nd^ millennium BCE.

Siwa Oasis and the Libyan region in general could be a region of prime importance for the understanding of date palm origins in North Africa and the history of gene flows with the Cretan date palm *Phoenix theophrasti*. More precisely, not only date palms from the current oasis could shed light on the history of this species, but also the uncultivated palms from the abandoned oasis. So far, research has focused mainly on cultivated germplasm, neglecting uncultivated date palms, even if local oasis communities have a use for them. Here we demonstrate that they can be of interest as, not only they can inform on past history, but they may represent untapped reservoir of diversity for future breeding programs.

### 4.5. Articulating the scales of ethnography and domestication over the long term

This study demonstrates that the combination of observation angles and methodologies on the cultivation of a crop, in particular perennial with a clonal/sexual reproduction, is essential. Ethnobotany hypothesized agricultural and classification practices with considerable effects on agrobiodiversity, made possible by local ways of categorizing living organisms (that result in the *de facto* presence of cultivars, ethnovarieties and local categories of date palms). Nevertheless, their occurrences are rare events (on the scale of one generation of farmers) and only molecular genetic analyses could verify those hypotheses. The other way around, geneticists assessing agrobiodiversity *de facto* consider each named type to be a cultivar, which only a long/careful ethnography could hypothesize to be, at least partially, untrue. Without the ethnobotanists, geneticists, by collecting a single sample per named type as they consider that they are cultivars, miss a huge amount of diversity and ways to understand it. The crucial step of sampling strategy cannot also be effectively implemented without a well-established understanding of local ways of categorizing (naming) the material being studied through long-term ethnobotanical field work.

In this study of the oasis of Siwa, we thus demonstrated that agrobiodiversity can be studied neither by a single genetic approach, nor by a single ethnographic approach. The combination of the two observation scales is necessary to uncover the phenomena and data necessary to test hypotheses. Indeed, these dialogues and close collaborations have resulted here in the discovery of hidden date palm diversity. Applying such an integrated approach to other oases may bring to light concealed date palm diversity elsewhere. Date palm diversity, whether wild, cultivated or abandoned thus still remains largely uncovered. Yet, having the right picture of the existing date palm diversity is crucial to the understanding of its origins, and critical when it comes to germplasm conservation.

## Supplementary materials

Table S1. *Phoenix* spp. accessions included in this paper. (Excel file)

Table S2. Microsatellite genotypes of 176 *Phoenix dactylifera* from Siwa region, nine *Phoenix theophrasti* and two *Phoenix reclinata*. (Excel file)

Table S3. Identity analysis on each pair of samples. (Excel file)

Table S4. Pairwise F_ST_ among various populations and subpopulations.

Table S5. Ancestry proportions in the Structure analysis (column) of the four original populations defined based on geographic origins of sample (row).

Figure S1. Pictures.

Figure S2. Principal Component Analysis of 347 *Phoenix dactylifera* accessions genotyped across 17 nuclear microsatellites.

Figure S3. Principal Component Analysis of *Phoenix* spp. accessions genotyped across 17 nuclear microsatellites.

Figure S4. Ancestry plot derived from Structure analysis of 356 accessions of *Phoenix dactylifera* and *Phoenix theophrasti* genotyped across 17 nuclear microsatellite loci.

Figure S5. Plots of (A) average maximum log likelihood over the 10 runs for the Structure analysis performed on 356 date palm and *Phoenix theophrasti* accessions (ancestry plot in Figure 5 and Figure S4) and (B) delta *K* from the same Structure analysis.

Figure S6. Allelic (left) and private allelic (right) richness. Calculated in two (A), three (B) and five (C) populations, using the rarefaction method (haploid sample size = 18, bootstrap number = 1,000).

Figure S7. Zoom on Siwa accessions only, from ancestry plot derived from 356 accessions of *Phoenix dactylifera* and *Phoenix theophrasti* genotyped across 17 nuclear microsatellite loci.

Figure S8. Distribution of the chlorotypes in *Phoenix dactylifera* and *Phoenix theophrasti* accessions.

## Acknowledgements

The present study was funded by BioDivMeX program (working-group Insularities)/MISTRALS, Mosaïque program (GDR 3355 INEE CNRS), and a grant from the New York University Abu Dhabi Research Institute on Biodiversity Genomics to Michael Purugganan. The authors are very grateful to the countless farmers of Siwa Oasis for having taken on such countless questions and for their unfailing hospitality.

## Data Archiving Statement

Genotyping data generated for this study are available in Table S2.

