## Supplementary material for "Integration of ethnobotany and population genetics uncovers the agrobiodiversity of date palms of Siwa Oasis (Egypt) and their importance to the evolutionary history of the species": Supp

**Table S1. *Phoenix* spp. accessions included in this paper.** (Excel file)

The field “Country” indicates the country of origin of the sample. For some cultivars, it may be different from the place of sampling, as they may be cultivated in a region, but known to originate from another one.

The field “Reference” indicates whether it is new data (from this paper, hence found in Table S2) or if it was sourced from a published study.

For Siwa named types, the local name of the dates is given (sometimes followed, after a /, by an alternative name or pronunciation), followed by the possible name of the palm if different [in brackets], and the possible name in local Arabic (in parentheses) (see Battesti, 2013 for explanation about the synonymy issue).

**Table S2. Microsatellite genotypes of 176 *Phoenix dactylifera* from Siwa region, nine *Phoenix theophrasti* and two *Phoenix reclinata*.** (Excel file)

CpM12 refers to the genotyping data of the chloroplastic minisatellite while the following columns reports genotyping data for the 17 nuclear microsatellites.

For nuclear microsatellites, the two alleles are separated by "/".

Missing data are indicated with NAs.

More information on the samples and the loci can be found in Table S1 and Table 2, respectively.

**Table S3. Identity analysis on each pair of samples.** (Excel file)

The number of nuclear microsatellite loci typed for each sample is reported (less than 17 indicates missing data), along with the number of matching and mismatching loci.

**Table S4. Pairwise F_ST_ among various populations and subpopulations.**

| (A) |  |  |  |  |  |  |
| --- | --- | --- | --- | --- | --- | --- |
|  | Eastern^+^ | *P. theophrasti* |  |  |  |  |
| *P. theophrasti* | 0.4029 |  |  |  |  |  |
| Western (Siwa included) ^+^ | 0.0875 | 0.3172 |  |  |  |  |
| (B) |  |  |  |  |  |  |
|  | Eastern | Siwa* | *P. theophrasti* |  |  |  |
| Siwa | 0.1193 |  |  |  |  |  |
| *P. theophrasti* | 0.4029 | 0.3257 |  |  |  |  |
| Western (Siwa excluded) ^+^ | 0.0843 | 0.0566 | 0.3346 |  |  |  |
| (C) |  |  |  |  |  |  |
|  | Eastern^+^ | Siwa Oasis^☨^ | Siwa uncultivated^☨^ | *P. theophrasti* |  |  |
| Siwa Oasis^☨^ | 0.1252 |  |  |  |  |  |
| Siwa uncultivated^☨^ | 0.1420 | 0.0547 |  |  |  |  |
| *P. theophrasti* | 0.4029 | 0.3517 | 0.2796 |  |  |  |
| Western (Siwa excluded) ^+^ | 0.0843 | 0.0690 | 0.0505 | 0.3346 |  |  |
| (D) |  |  |  |  |  |  |
|  | Eastern^+^ | Siwa cultivated^•^ | Siwa úšik^•^ | Siwa uncultivated^☨^ | *P. theophrasti* |  |
| Siwa cultivated^•^ | 0.1332 |  |  |  |  |  |
| Siwa *úšik* ^•^ | 0.1138 | 0.0070 |  |  |  |  |
| Siwa uncultivated^☨^ | 0.1420 | 0.0673 | 0.0316 |  |  |  |
| *P. theophrasti* | 0.4029 | 0.3694 | 0.3344 | 0.2796 |  |  |
| Western (Siwa excluded) ^+^ | 0.0843 | 0.0781 | 0.0534 | 0.0505 | 0.3346 |  |
| (E) |  |  |  |  |  |  |
|  | Eastern^+^ | Siwa cultivated^•^ | Siwa úšik #1^∇^ | Siwa úšik #2^∇^ | Siwa uncultivated^☨^ | *P. theophrasti* |
| Siwa cultivated^•^ | 0.1332 |  |  |  |  |  |
| Siwa úšik #1^∇^ | 0.1192 | 0.0026 |  |  |  |  |
| Siwa úšik #2^∇^ | 0.111 | 0.0138 | 0.0028 |  |  |  |
| Siwa uncultivated^☨^ | 0.142 | 0.0673 | 0.0392 | 0.0216 |  |  |
| *P. theophrasti* | 0.4029 | 0.3694 | 0.3301 | 0.3566 | 0.2796 |  |
| Western (Siwa excluded) ^+^ | 0.0843 | 0.0781 | 0.0599 | 0.047 | 0.0505 | 0.3346 |

^+^ *Eastern* and *Western* accessions refer to date palm accessions from North Africa and the Middle East up to Pakistan, respectively.

* *Siwa* refers to the 128 unique genotypes sampled in the oasis (named types, úšik #1 and #2) and in the surrounding desert (uncultivated date palms)

☨ *Siwa Oasis* and *Siwa uncultivated* refer to date palm accessions sampled in the oasis, and uncultivated accessions from the surrounding desert, respectively.

^•^ *Siwa cultivated* refers to named types from the oasis while *Siwa úšik* refers to both úšik #1 and #2 accessions in Siwa Oasis, respectively.

^∇^ Siwa úšik #1 and #2 refer to accidental seedlings growing respectively #1 in gardens, and #2 at the edge of gardens or palm groves.

**Table S5. Ancestry proportions in the Structure analysis (column) of the four original populations defined based on geographic origins of sample (row).**

| **Population origin** | **Average ancestry at *K* = 4** | | | |
| --- | --- | --- | --- | --- |
|  | **Eastern** | **Western** | **Siwa** | ***P. theophrasti*** |
| **Eastern** | **97.50** | 1.34 | 0.93 | 0.19 |
| **Western** | 6.48 | **80.95** | 11.85 | 0.71 |
| **Siwa** | 2.73 | 7.66 | **88.38** | 1.24 |
| ***P. theophrasti*** | 0.47 | 3.46 | 0.62 | **95.43** |

With *K* = 4, we can identify four clusters that mostly correspond to the four populations as defined based on their geographic origin.

Figure S1. Pictures.

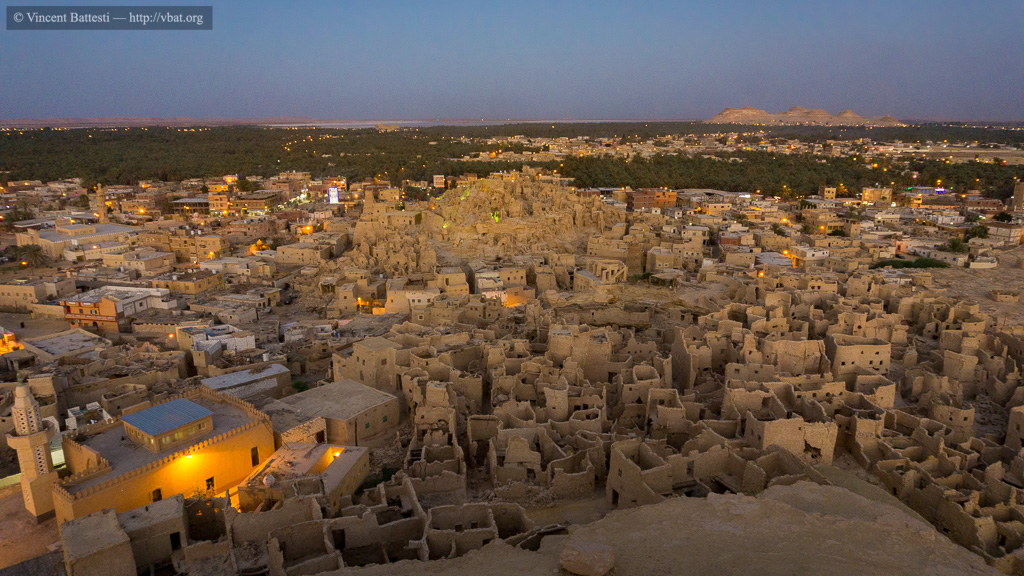

(A) View of the eastern part of Siwa Oasis (Egypt) from its urban center at nightfall. In the foreground stands out the ancient city (Shali) built in salt clay, and in the background stands out the palm grove, and just before one of the salt lakes the inselberg of Aghurmi where the oracle temple of Amon is erected. September 27, 2017 (6:30 pm), Vincent Battesti

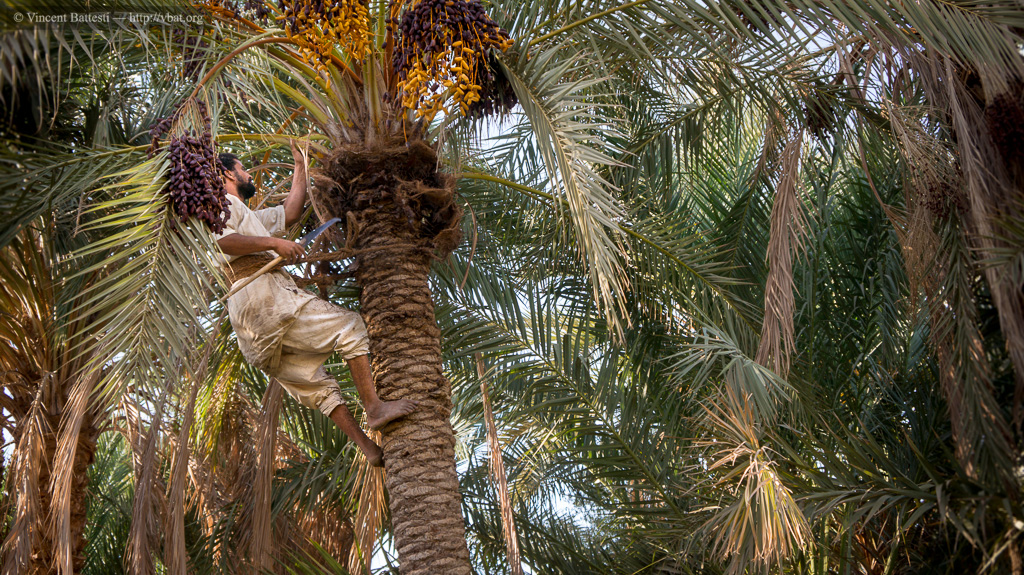

(B). In a garden in the Siwa palm grove (in the Tanɣāzi area), one of the many Isiwan farmers who work on harvesting dates, here with his emblematic local sickle helped (this is not always the case) by a belt made of date tree fibers. These úšik dates are rather intended for fodder consumption. November 11, 2014 (3:30 pm), Vincent Battesti

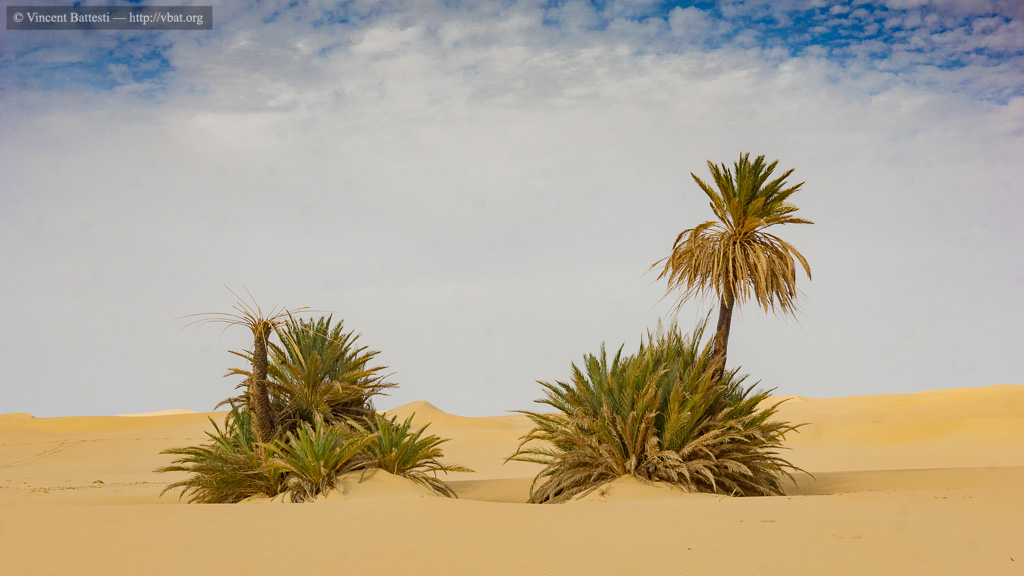

(C). Two examples of uncultivated date palms in an area called Labbaq south of Siwa Oasis (in the early Great Sand Sea). These two date palms are part of our sample (3467 and 3468) which have been found to be the same genotype (off-shoots of off-shoots that have separated over centuries from each other by 7 to 10 m). September 25, 2017 (4:00 pm), Vincent Battesti.

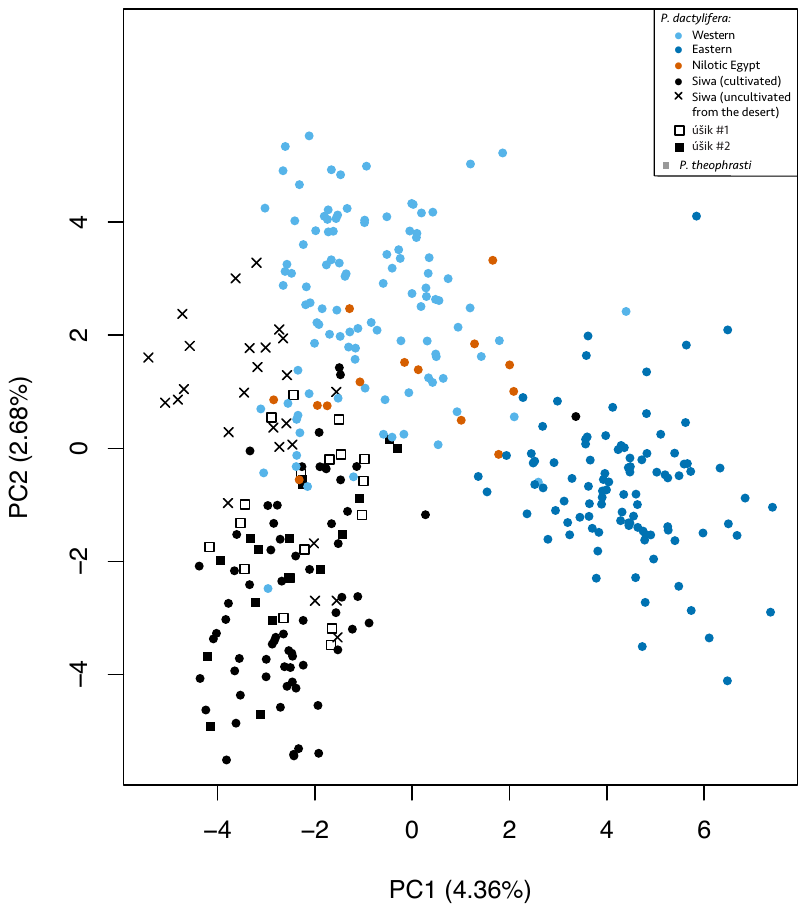

Figure S2. Principal Component Analysis of 347 *Phoenix* *dactylifera* accessions genotyped across 17 nuclear microsatellites.

This is the same Figure as in Figure 5B except that seedlings sampled in the garden (úšik #1) and on the border of the gardens or palm groves (úšik #2) are shown with square. We can see that they cluster within the Siwa cluster, with oasis and desert date palms.

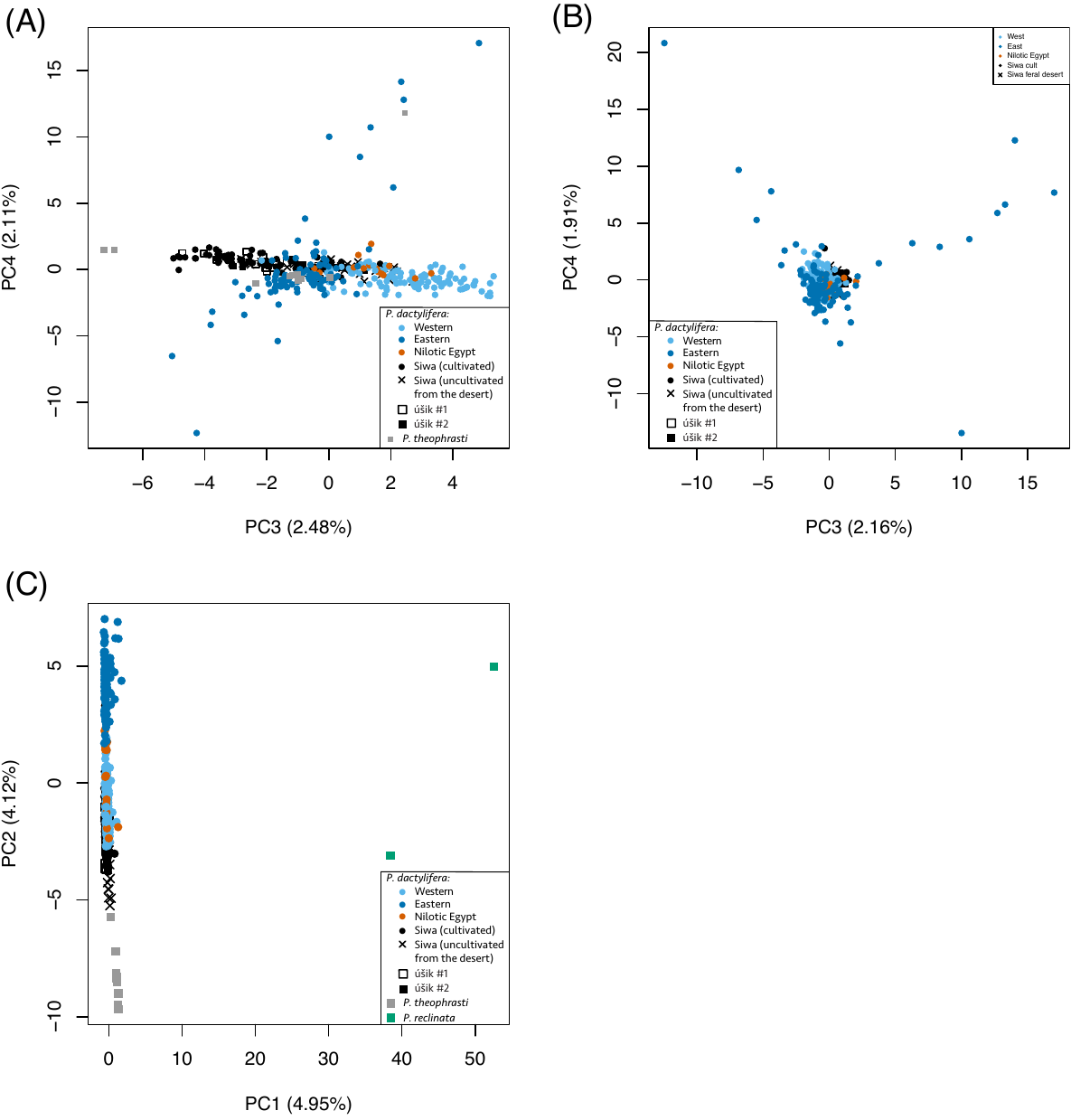

**Figure S3. Principal Component Analysis of *Phoenix* spp. accessions genotyped across 17 nuclear microsatellites.**

(A) PCA on nine *Phoenix theophrasti* and 347 *Phoenix dactylifera* (variance explained in parentheses). Plot of PC1 and 2 can be found in Figure 5A.

(B) PCA on 347 *Phoenix dactylifera* (variance explained in parentheses). Plot of PC1 and 2 can be found in Figure 5B. Same legend as in (A).

(C) PCA on 2 *Phoenix reclinata,* nine *Phoenix theophrasti* and 347 *Phoenix dactylifera*.

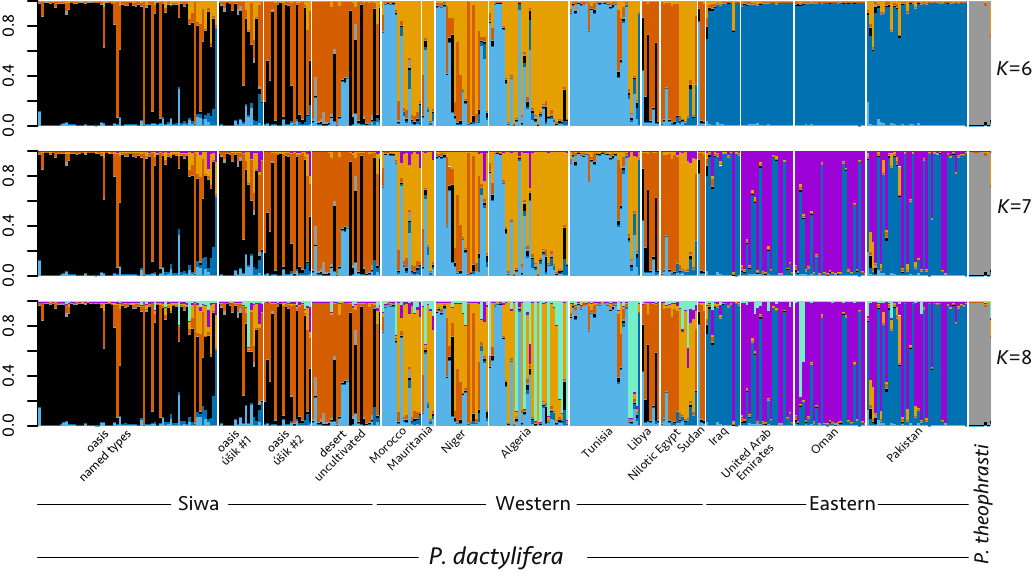

Figure S4. Ancestry plot derived from Structure analysis of 356 accessions of *Phoenix dactylifera* and *Phoenix theophrasti* genotyped across 17 nuclear microsatellite loci.

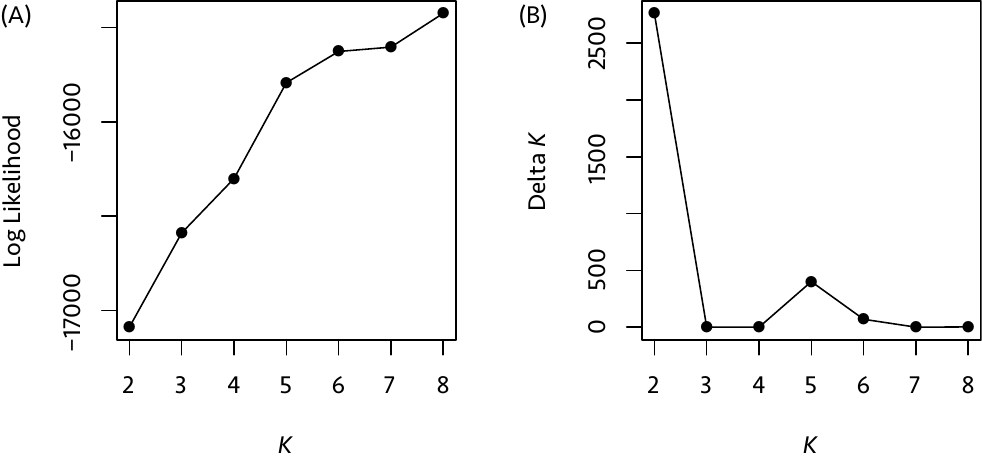

Figure S5. Plots of (A) average maximum log likelihood over the 10 runs for the Structure analysis performed on 356 date palm and *Phoenix theophrasti* accession (ancestry plot in Figure 5 and Figure S4) and (B) delta *K* from the same Structure analysis.

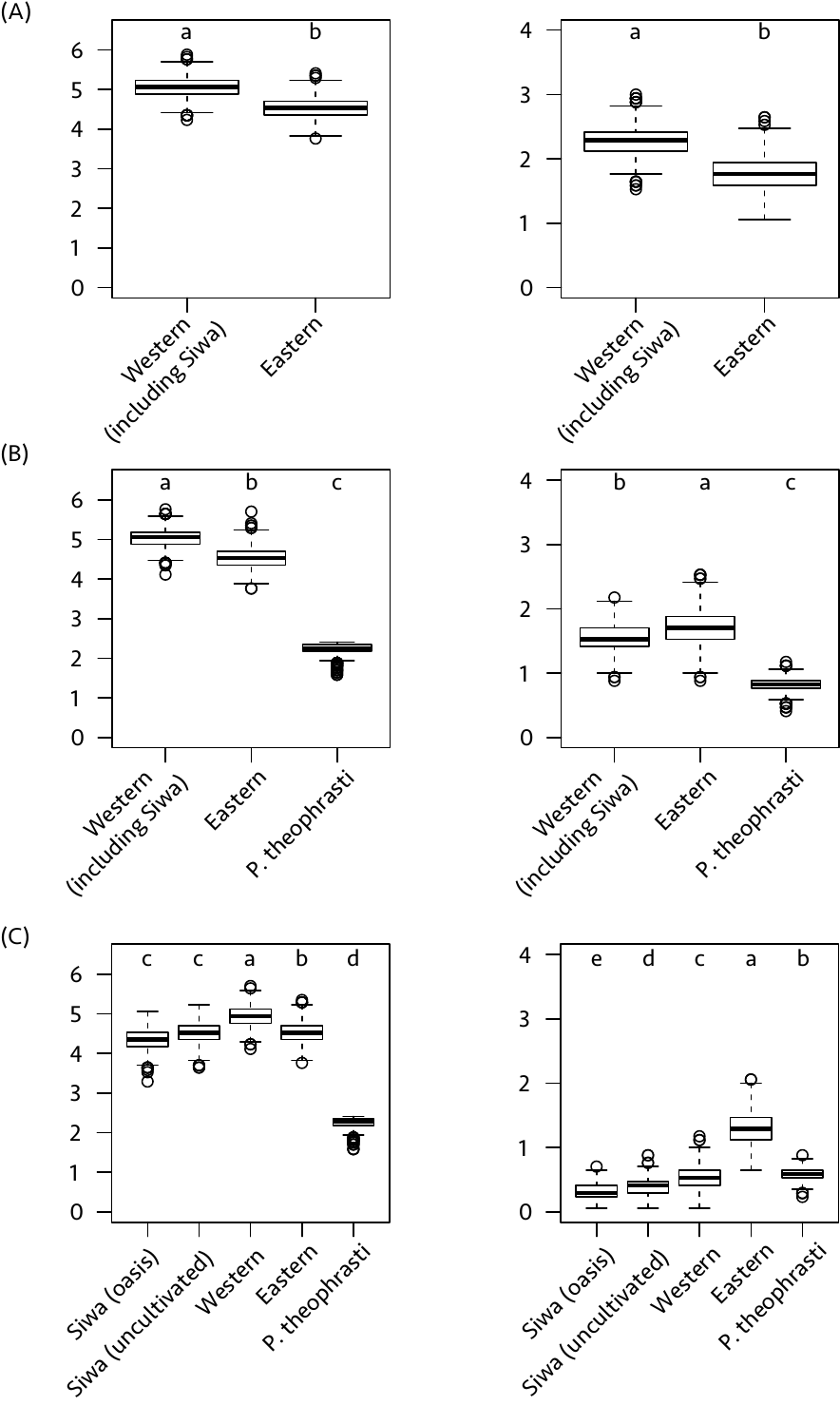

Figure S6. Allelic (left) and private allelic (right) richness. Calculated in two (A), three (B) and five (C) populations, using the rarefaction method (haploid sample size = 18, bootstrap number = 1,000).

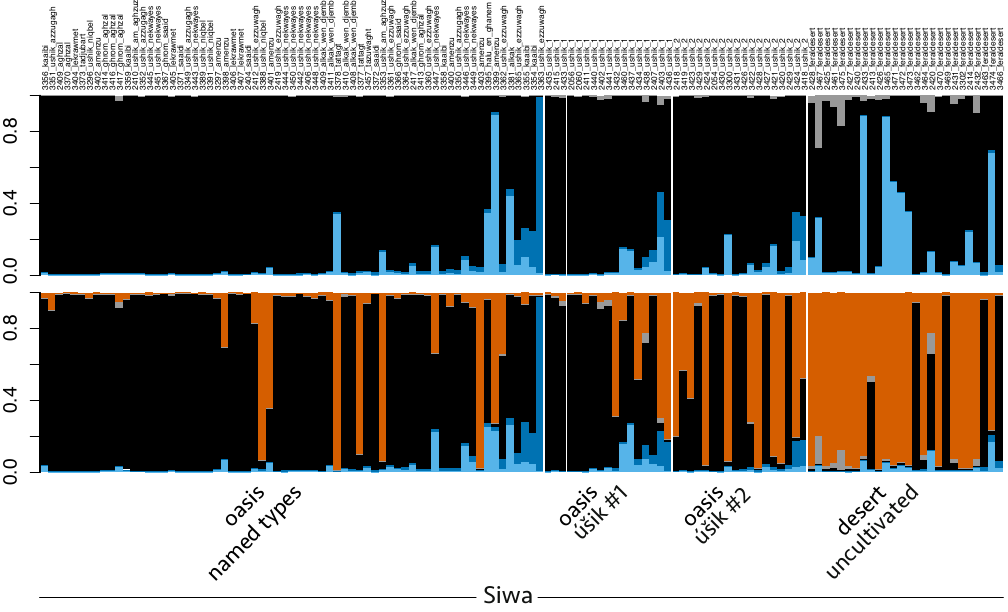

Figure S7. Zoom on Siwa accessions only, from ancestry plot derived from 356 accessions of *Phoenix dactylifera* and *Phoenix theophrasti* genotyped across 17 nuclear microsatellite loci.

The plot with all samples can be seen in Figure 5C and Figure S3. Top: *K* = 4; bottom: *K* = 5.

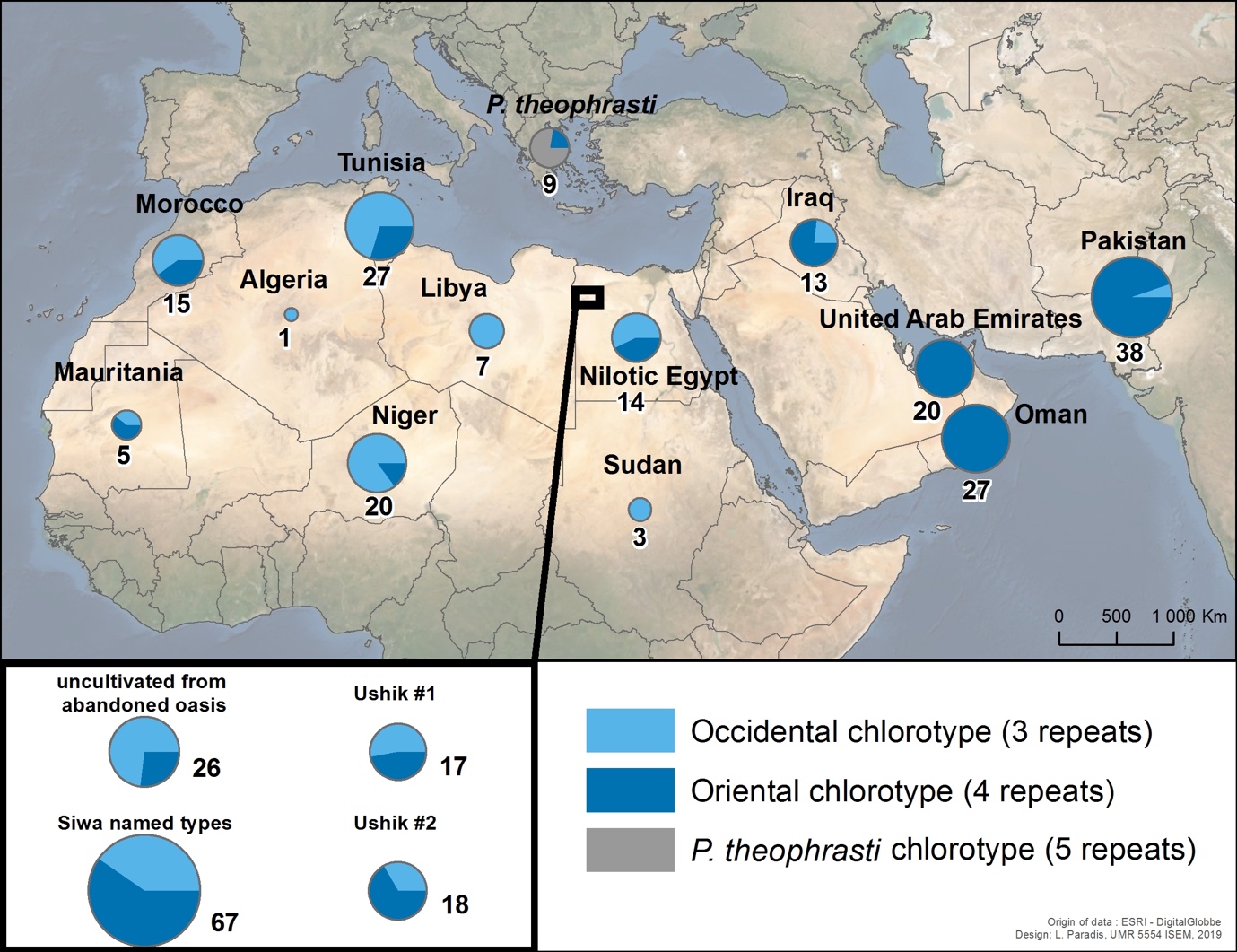

Figure S8. Distribution of the chlorotypes in *Phoenix dactylifera* and *Phoenix theophrasti* accessions.
